# Investigating the prognostic value of *KCNN4* gene expression in human pancreatic adenocarcinoma by bioinformatic analysis

**DOI:** 10.1101/2025.04.26.650766

**Authors:** Syed Sajidul Islam, Md. Mehedi Hasan Tushar, Bishajit Sarkar, Ripa Moni, Umme Salma Zohora, Mohammad Shahedur Rahman

**Author notes:** Corresponding authors: Umme Salma Zohora, Mohammad Shahedur Rahman Email of the corresponding authors.

## Abstract

Finding accurate biomarkers for early detection and prognosis is critical because pancreatic adenocarcinoma (PAAD) is a deadly cancer with a poor prognosis. Though its function in PAAD is still unknown, the gene *KCNN4*, which codes for a calcium-activated potassium channel, has been linked to a number of malignancies. In this work, the cancer studies of the CPTAC, TCGA, and GTEx projects were targeted using a thorough database mining technique. Several analytical methods were used to examine full-scale molecular, expression, and prognostic profiles of the *KCNN4* gene in pancreatic adenocarcinoma tissues. It was discovered that the majority of the cancerous tissue had different levels of mRNA and protein expression for this gene. The results of comparative immunohistochemistry also revealed that normal pancreatic tissues had reduced expression of the KCNN4 protein. An analysis overview of copy-number alterations and *KCNN4* mutations in PAAD samples was also conducted. Furthermore, there was a correlation found between the expression of *KCNN4* and both overall and disease-free survival in pancreatic adenocarcinoma. In pancreatic adenocarcinoma tissue, co-expressed genes of *KCNN4* were found to be involved in cell maintenance-related functions, according to gene coexpression, additional ontology, and pathway analysis. When it comes to cancer diagnosis and treatment, the experimental findings of this study should aid in integrating *KCNN4* into clinical applications.

## 1. Introduction

### 1.1 The global prevalence of cancer

Cancer is a disease characterized by abnormal metabolism and signaling, which allows transformed cells to divide uncontrollably and survive. Numerous chemicals, elements, and circumstances have been identified as fundamental contributors to the onset and development of the disease (Upadhyay, 2021). One of the most deadly groups of diseases affecting humans, it causes millions of deaths worldwide every year and exhibits a wide range of characteristic clinical features. These disease groups include over a hundred genetically distinct conditions that have certain similarities in their molecular mechanisms and metabolic changes (Lambert et al., 2017; Vander Heiden & DeBerardinis, 2017).

Cancer is a growing global health concern that presents significant diagnostic and therapeutic challenges. According to the World Health Organization (WHO), cancer will be the leading cause of death worldwide in 2020, accounting for approximately 10 million deaths (de Martel et al., 2020; Ferlay et al., 2020; World Health Organization, 2023). The most prevalent kinds are stomach, colorectal, lung, prostate, and skin (non-melanoma) cancers (Siegel et al., 2018).

The growing rate of cancer is caused by several factors, such as changing lifestyles, aging populations, and environmental exposures. Growing older is a significant risk factor because it increases the chance of getting cancer because genetic mutations accumulate over time (Islami et al., 2018). Additionally, a person’s risk of developing cancer is greatly increased by lifestyle choices like smoking, drinking alcohol, eating poorly, and not exercising (Torre et al., 2016).

Pancreatic cancer is the 12th most frequent cancer and ranks sixth in terms of fatality rates worldwide in both sexes, according to GLOBOCAN 2022 estimates. According to reports, North America, Europe, and Australia-New Zealand have the highest occurrence rates of cancer (Bray et al., 2024). From 1991 onward, pancreatic cancer incidence and mortality increased in practically every country in the world (Ilic & Ilic, 2022, 2024). Conversely, only Canada and Mexico reported favorable trends in pancreatic cancer mortality. According to a comparative mortality study conducted across 28 European Union countries, pancreatic cancer is expected to overtake breast cancer by 2025 (i.e., there will be a 25% increase in pancreatic cancer deaths over breast cancer deaths), placing it on the third position of the most frequent cause of mortality from cancer by that time (after colorectal and lung cancer) (Ferlay et al., 2016).

**Fig 1:**
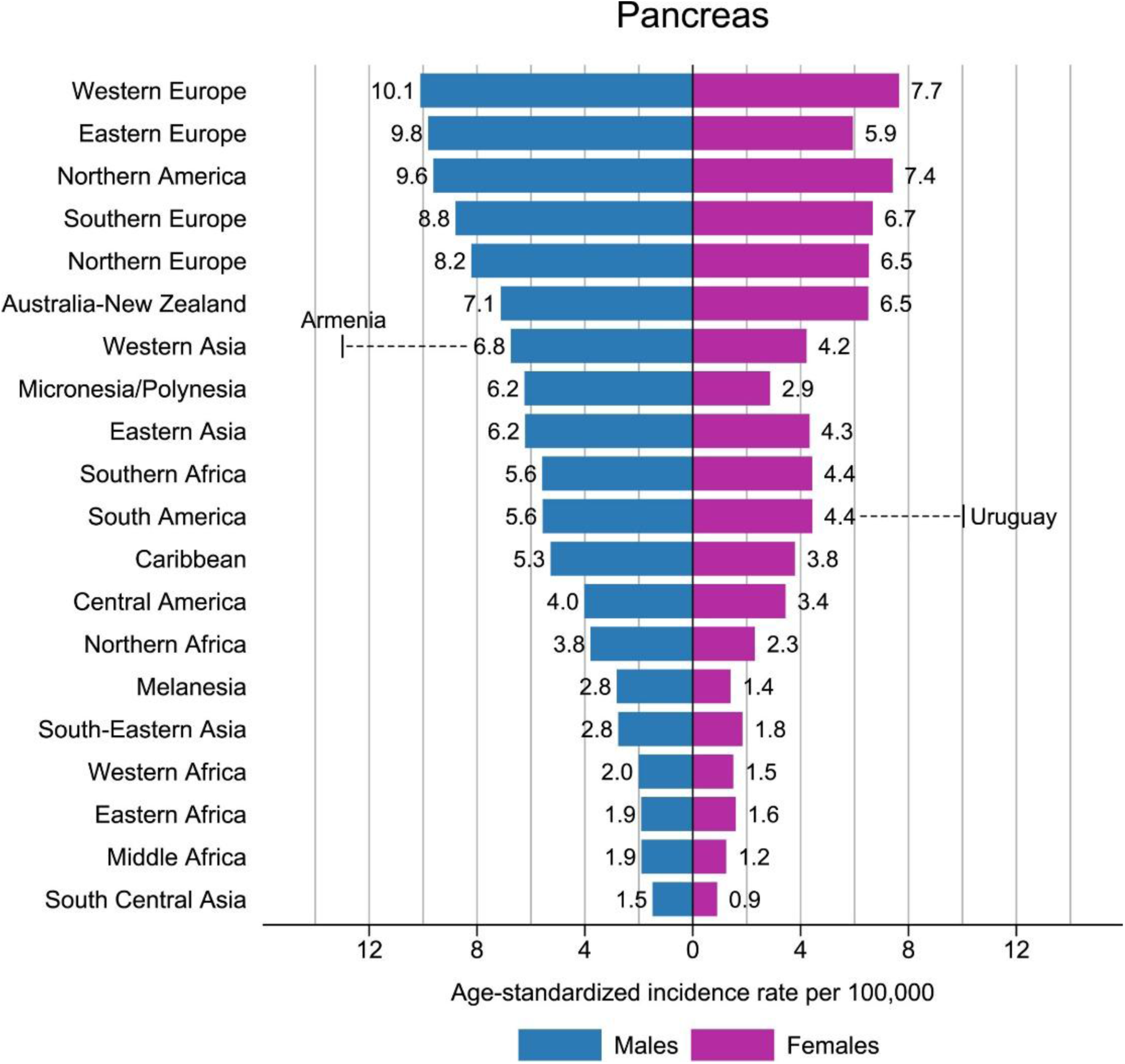
The age-standardized rate of pancreatic cancer incidence by sex and the rates specific to a certain region are shown in this column graph. The world age-standardized rate for males is shown in descending order, with the highest national rates for both males and females superimposed. (Source: GLOBOCAN 2022) (Bray et al., 2024)

Early cancer diagnosis is essential for better survival rates and efficient treatment, but it is still very difficult. Screening programs and other early detection techniques vary in their accessibility and efficacy. For instance, it has been demonstrated that mammography for breast cancer and colonoscopy for colorectal cancer lower death rates; however, there are differences in the accessibility of these tests, especially in low- and middle-income nations (Jemal et al., 2006).

Furthermore, some cancers—like pancreatic cancer—don’t have good early detection techniques, which frequently leads to late-stage diagnoses and dismal prognoses (Rahib et al., 2014).

### 1.2 Biological functions of the *KCNN4* gene

The calcium-activated potassium channel protein with intermediate conductance is encoded by the KCNN4 gene, also referred to as KCa3.1. Numerous tissues, including hematopoietic cells, epithelial cells, and the vascular endothelium, exhibit high levels of expression for this channel. It is essential for many physiological functions, most notably the control of intracellular calcium signaling and membrane potential. The control of potassium ion flow in response to intracellular calcium levels is the principal biological role of the KCNN4 channel. Membrane hyperpolarization results from the opening of the KCNN4 channel, which permits potassium ions to leave the cell in response to an increase in intracellular calcium. In turn, this hyperpolarization affects a number of cellular processes, including the control of cell volume, proliferation, and migration (Feske et al., 2015; Joiner et al., 1997; Wei et al., 2005).s

KCNN4 channels are essential for maintaining ionic homeostasis because they allow the efflux of K+ ions in response to increases in intracellular Ca2+ concentrations. Numerous physiological processes, including migration, proliferation, and regulation of cell volume, depend on this mechanism to function appropriately (Faber & Sah, 2003). For example, in erythrocytes, KCNN4 mediates K+ efflux, which is essential for these cells to survive under osmotic stress and helps maintain cell volume and shape (Hoffman et al., 2003).

Activation and proliferation of T lymphocytes in the immune system are regulated by the expression of KCNN4 channels. The channel is essential to immune responses because its activity maintains the Ca2+ signaling required for T-cell activation and cytokine production (Wulff et al., 2000).

### 1.3 *KCNN4* in the oncogenic processes

Current research has demonstrated *KCNN4*’s role in a number of oncogenic processes. Numerous cancer types have been linked to the channel’s involvement in the invasion, migration, and multiplication of cancer cells. It seems essential to the survival and aggressive behavior of cancer cells for it to keep the ionic balance within them (George Chandy et al., 2004).

In pancreatic cancer, *KCNN4* contributes to the aggressive characteristics of cancer cells. By preserving the ionic homeostasis necessary for these functions, the channel’s activity promotes the continuous growth and survival of pancreatic cancer cells (Jäger et al., 2004). Furthermore, KCNN4 affects how cancer cells interact with their surroundings, which promotes the growth and metastasis of tumors (Jensen et al., 2002; Ohya & Kito, 2018).

### 1.4 Objectives of the study

i. To investigate the potential oncogenic roles and clinical implications of *KCNN4* gene expression in human pancreatic adenocarcinoma (PAAD).
ii. To evaluate their expression pattern, methylation, and mutation status as well as their overall impact on cancer patients’ survival.
iii. To guide the upcoming clinical research on *KCNN4* and its byproducts for the purpose of developing new cancer treatments and diagnostic tools.

## 2. Materials and Methods

### 2.1 Analysis of expression of *KCNN4* in various normal tissue and cancerous tissue types

Two different online based tools, i.e., GEPIA2 (http://gepia2.cancer-pku.cn/#index), and GENT2 (http://gent2.appex.kr/), were utilized to investigate the different expressions pattern of KCNN4 across a variety of normal calls and cancer cells (Park et al., 2019, p. 2; Tang et al., 2017). A tool for various transcriptional analyses, such as differential-expression analysis and correlation in a variety of cancerous and normal tissues, is GEPIA2, which is built-in RNA-sequencing expression data from the GTEx and the TCGA projects, comprising 8,587 normal samples and 9,736 tumors. (Tang et al., 2017). During the analysis with the GEPIA2 server, default values for the parameters were used. On the other hand, GENT2 offers expression pattern analysis across 72 different tissues and comprises over 68,000 samples, making it a more valuable platform for prognostic analysis of targeted genes (Park et al., 2019).

### 2.2 *KCNN4* expression pattern optimization in normal and pancreatic adenocarcinoma (PAAD) tissue

Three web-based tools were utilized in this method, i.e., GEPIA2 server, UALCAN server (https://ualcan.path.uab.edu), and the HPA server (www.proteinatlas.org) (Chandrashekar et al., 2017; Pontén et al., 2008; Tang et al., 2017). Using a combination of TCGA and GTEx datasets, the GEPIA2 web-server was utilized to compare the differences in *KCNN4* gene expression between the normal tissues and those affected by pancreatic adenocarcinoma. An integrated, interactive resource is the UALCAN web server which is built on several widely accessible OMICS databases. It is useful for a variety of tasks, such as facilitating the interpretation of expression profiles, demonstrating patient survival characteristics, and controlling gene expression through epigenetic regulation. As with the prior step, we used this web server to determine our target genes’ expression profile in the normal tissue and pancreatic adenocarcinoma (PAAD) tissue. Through the combination of several sets of omics technologies, like transcriptomics, mass spectrometry (MS) based proteomics, antibody-based imaging, and systems biology, the Human Protein Atlas (HPA) facilitates the investigation of the human proteome (Pontén et al., 2008). Using the HPA server, a graphic assessment of the histological samples (HPA053841) among the normal and pancreatic adenocarcinoma (PAAD) tissue was produced. In this step, every experiment was conducted using the default parameters, with a p value of less than 0.05 being considered important.

### 2.3 Determining the pattern of methylation in DNA and *KCNN4* promoter in pancreatic adenocarcinoma (PAAD)

UALCAN web server was utilized to search for variations throughout the *KCNN4* gene’s promoter-methylation pattern in both normal and pancreatic tumor tissues. After that, GDC TCGA Pancreatic Cancer (PAAD) samples (n=223) had been selected as the basis sample set, and using the UCSC-Xena web browser, the gene’s DNA methylation-pattern in PAAD tissues was determined (Chandrashekar et al., 2017; Goldman et al., 2017). We selected genomic data type of the KCNN4 gene utilizing gene expression, copy number, and somatic mutation dataset, while we left the other experiments parameters at their initial settings. Web-based tools for clinical and phenotypic annotation of a desired gene are provided by the UCSC Xena platform. With the addition of over 1500 datasets covering 50 distinct cancer types, this multi-omics database integration is enhanced. Along with gene expression, copy number, and somatic mutation analysis, the neoplasm histologic grade, disease type, and age at initial pathogenesis diagnosis was also analyzed in PAAD samples.

### 2.4 Analyzing the mutations in *KCNN4* and variations in copy numbers in pancreatic adenocarcinoma samples

The KCNN4 gene’s mutations and copy-number alterations were computed using the cBioPortal (www.cbioportal.org) web-based server (J. Gao et al, 2013, 2018; Jiao et al., 2019). It is an open access, web-based translational research platform that helps reveal the genomic and epigenomic differences associated with a given gene and allows for multidirectional analysis of that gene. Over 3,609 samples or 3,569 patients were examined in 13 different pancreatic cancer studies, and the server chose samples from among those deposited by the MSK, TCGA, QCMG, and CPTAC to look into copy-number alterations and mutations.

### 2.5 Investigation of the associations among *KCNN4* expression and clinical features of patients with pancreatic cancer

Utilizing the samples of TCGA supplied by the UALCAN web-based server, the associations among the *KCNN4* gene expression profile and various types of clinical characteristics of patients with pancreatic adenocarcinoma (PAAD) was ascertained (Chandrashekar et al., 2017). Different expression patterns were analyzed and recorded based on the specific cancer stages, age, race, gender, smoking history, histology of tumor, nodal-metastasis status, and presence of T53 mutation of the patient. The servers’ default settings were used for the experiment, and a p value of under 0.05 was used to classify predictions as statistically important.

### 2.6 Evaluating the association between pancreatic cancer patients’ probable survival trend and *KCNN4* expression

The Kaplan-Meier plotter online server (www.kmplot.com/analysis/) was utilized to establish the correlation among *KCNN4* expression and the survival-rate of pancreatic cancer patients (Nagy et al., 2018). More than 60,000 genes can be evaluated by this tool, including those that are involved in the expression of messenger RNA and microRNA in twenty one distinct types of cancer.

Various patterns of survival, such as overall survival (OS), and disease free survival (DFS) of pancreatic cancer patients in response to KCNN4 expression, were retrieved by drawing Kaplan-Meier plots while keeping other parameters default, and the analysis was conducted on 1237 patients. Thereafter, various *KCNN4* gene translational data related to pancreatic adenocarcinoma patients’ survival were also obtained from the PrognoScan (Mizuno et al., 2009) web database.

### 2.7 identifying different genes similar to *KCNN4* in pancreatic adenocarcinoma (PAAD)

The mining of different genes that are fairly comparable with KCNN4 was achieved by two distinct web servers, such as the GEPIA2, and UCSC-Xena web-server. Initially, the *KCNN4* gene was looked up in the GEPIA2 online database (Expression Analysis) to identify potential genes that are similar. According to related gene-clusters, the server produces Pearson correlation coefficient (PCC) values.

Thereafter, the similar genes were identified with PCC scores. Furthermore, the GEPIA2 web-based server was utilized to decipher and evaluate the relationship between the genes that were identified as very similar to KCNN4 in the preceding step and the KCNN4 gene. When presenting the relationship between the two chosen genes, the GEPIA2 server generated the p-values; any p-value lower than 0.05 was considered to be statistically noteworthy. Lastly, the gene-expression patterns of the 2 chosen genes in PAAD patients were predicted using the UCSC-Xena web browser.

### 2.8 Interpretation of ontology terms and pathways associated with KCNN4 and its functionally related genes

The Enrichr server (https://maayanlab.cloud/Enrichr/) was used to find the cell signaling pathways and gene ontology (GO) of KCNN4 and potentially related genes (E. Y. Chen et al., 2013). This platform utilizes enrichment analysis to infer information about a particular set of genes through comparison with multiple datasets of genomics that represent previous biological knowledge. Here, we employed the *KCNN4* gene and every potential gene from the preceding step that is similar to interpret various GO (gene ontology) terms, i.e., GO molecular function, GO biological processes, and GO cellular components. Subsequently, the Enrichr web-based server was utilized to extract multiple signaling pathways from the BioPlanet 2019, Reactome-2016, and KEGG 2019 Human databases.

## 3. Results and Discussion

### 3.1 Expression profile of *KCNN4* in a range of cancerous and normal tissues

The analysis of the GENT2 servers have detected a high expression of *KCNN4* gene in Pancreatic Adenocarcinoma (PAAD) tissues, with a pronounced presence of additional cancer types, such as Skin Cutaneous Melanoma (SKCM), Bladder Carcinoma (BLCA), Prostate Adenocarcinoma (PRAD), Rectum Adenocarcinoma (READ), Colon Adenocarcinoma (COAD), Cervical Squamous Cell Carcinoma and Endocervical Adenocarcinoma (CESC) and so on. On the contrary, KCNN4 was found to be under-expressed in Adenoid Cystic Carcinoma (ACC) and Cholangiocarcinoma (CHOL) tumor tissues based on our findings. These results were displayed through examination of the *KCNN4* gene expression patterns in the GEPIA2 (Figure 5) and GENT2 servers (Figure 6). Further comparisons between various cancer tissues and their corresponding normal counterparts revealed a higher expression of *KCNN4*.

**Fig 5:**
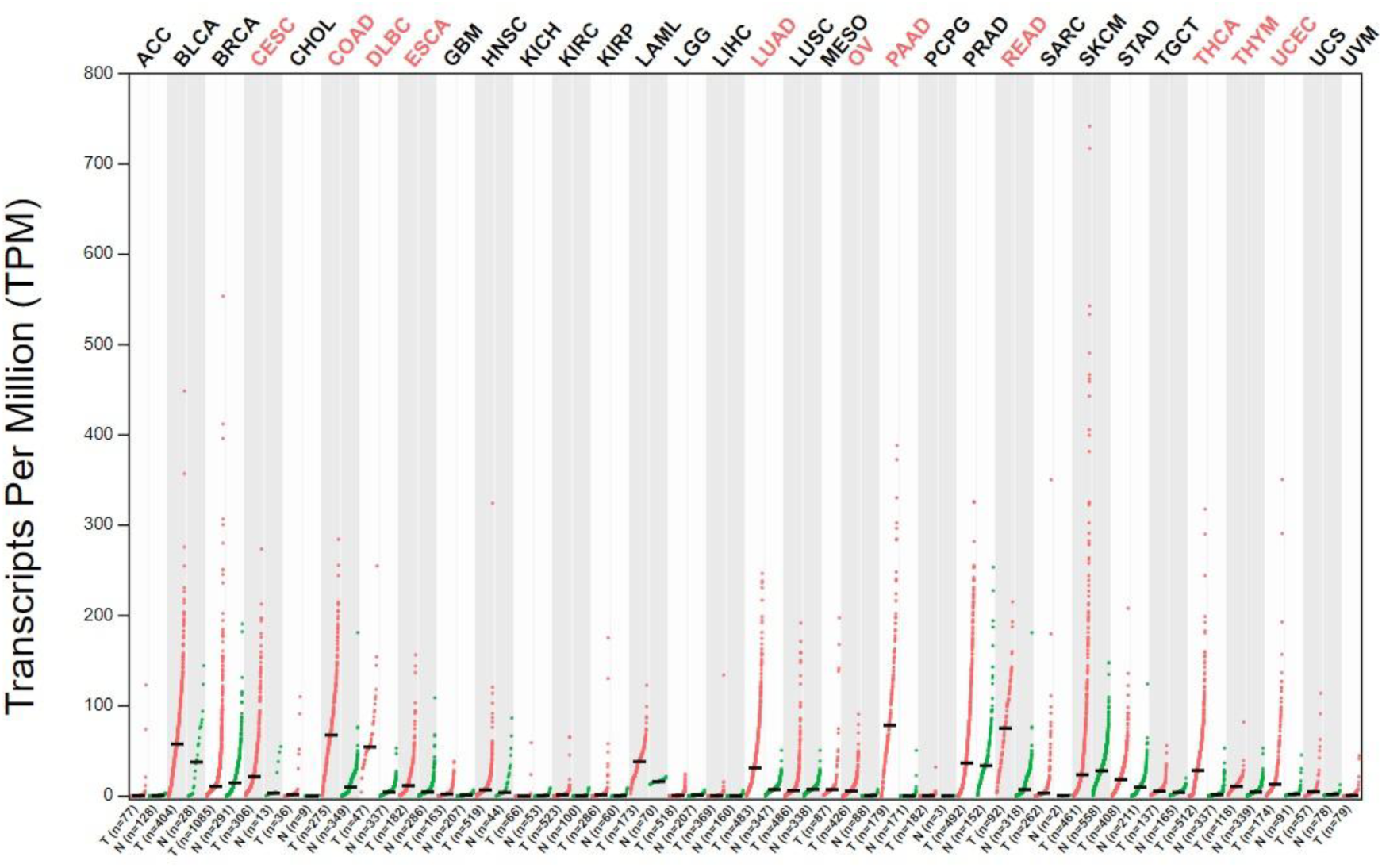
The GEPIA2 web server showed the overexpression of *KCNN4* (red) in a variety of tumor tissues.

**Fig 6:**
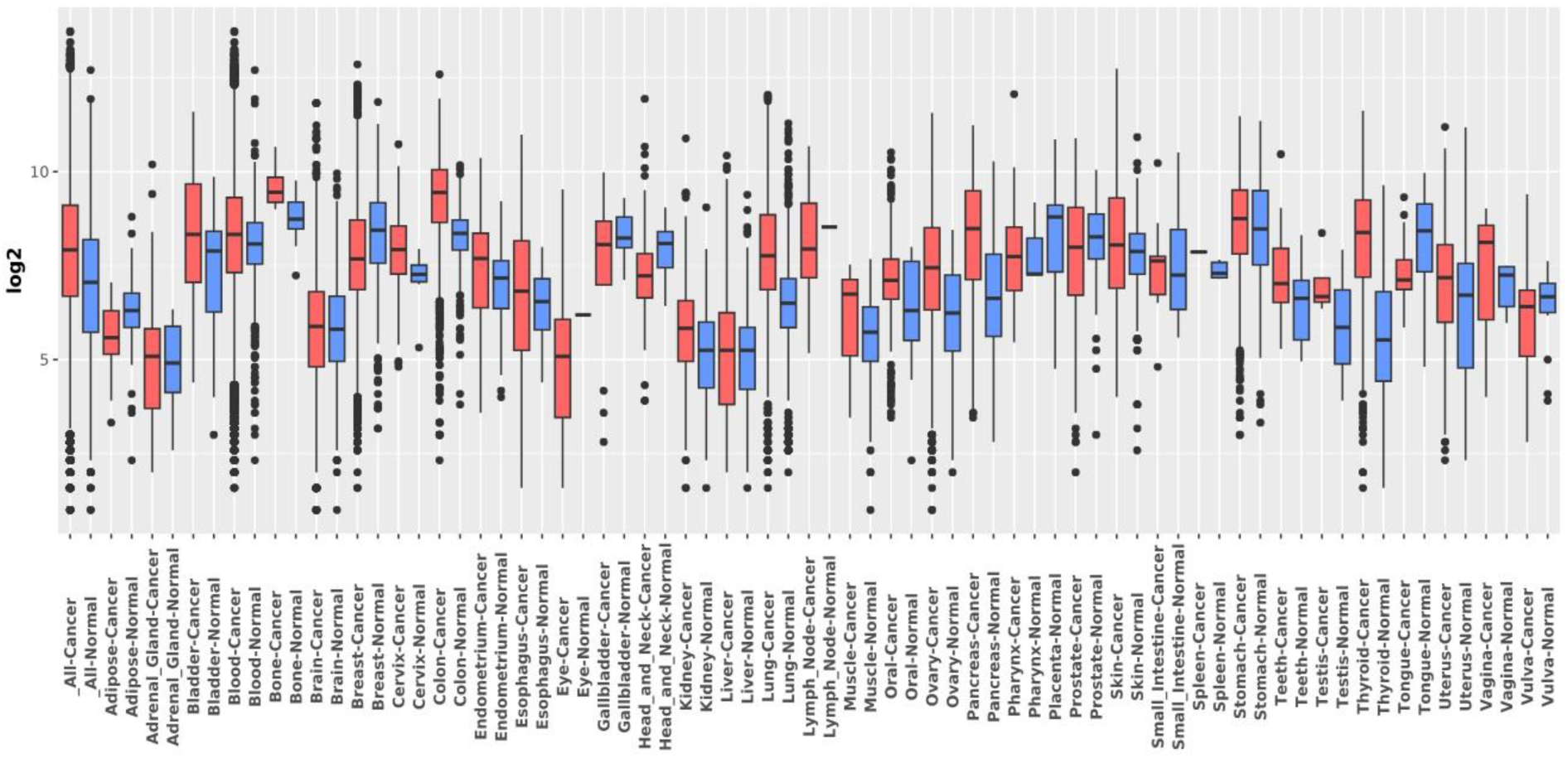
Based on the log 2-fold change in the GENT2 server, the boxplot interpretation illustrates the differences in KCNN4 expression across various cancer types.

### 3.2 *KCNN4* expression analysis in cancerous and normal pancreatic tissues

A similar investigation using the GEPIA2 server revealed results similar to the previous step, with the breast tumor samples (n = 179) exhibiting higher levels of KCNN4 expression and the median value of log2 (TPM+1) is converging around 6, in contrast to approximately 0.5 for the samples of normal pancreatic tissues (n= 171) (Figure 7). Once more, the analytical report on the TCGA samples obtained from the UALCAN web server indicated that breast tumor tissues (n=178) had higher levels of KCNN4 expression than normal tissues (n=4) (Figure 8). Subsequently, an analysis was conducted using the HPA server to compare the expressions of *KCNN4* protein levels in normal and cancerous pancreatic tissues by immunohistochemistry (IHC). The results showed that normal tissues exhibited no staining and a negative intensity of the immunohistochemistry test, while pancreatic cancer tissue had a weak assay intensity and no staining (Figures 9a and 9b). These assumptions reinforced the results of earlier investigations that indicated pancreatic cancer tissues expressed *KCNN4* slightly, both at the mRNA and protein levels, compared to the normal tissues.

**Fig 7:**
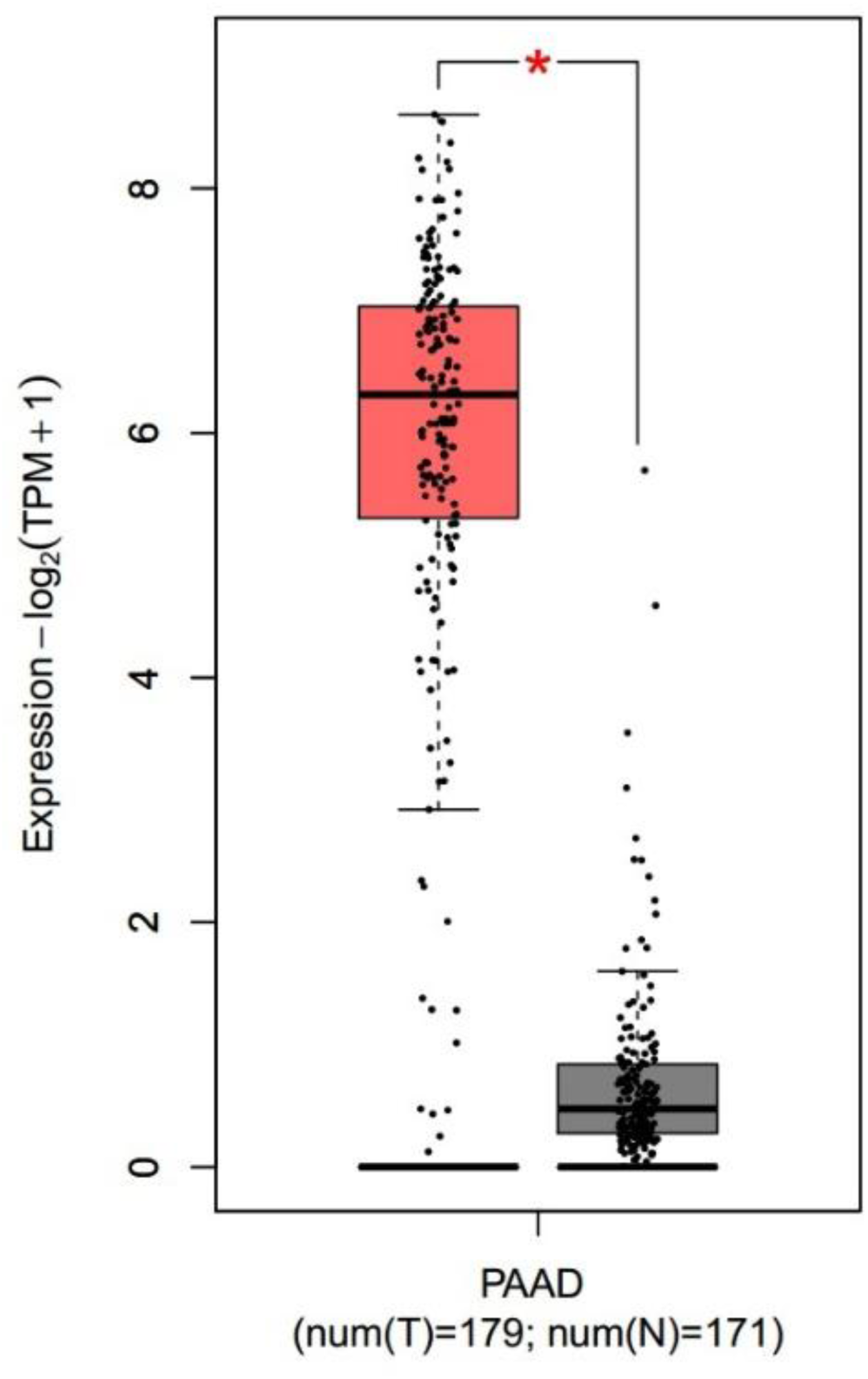
Investigation in the GEPIA2 server revealed that pancreatic adenocarcinoma (PAAD) samples had higher KCNN4 expression than normal breast tissues (the red box indicates PAAD samples, the black box indicates adjacent normal tissue samples)

**Fig 8:**
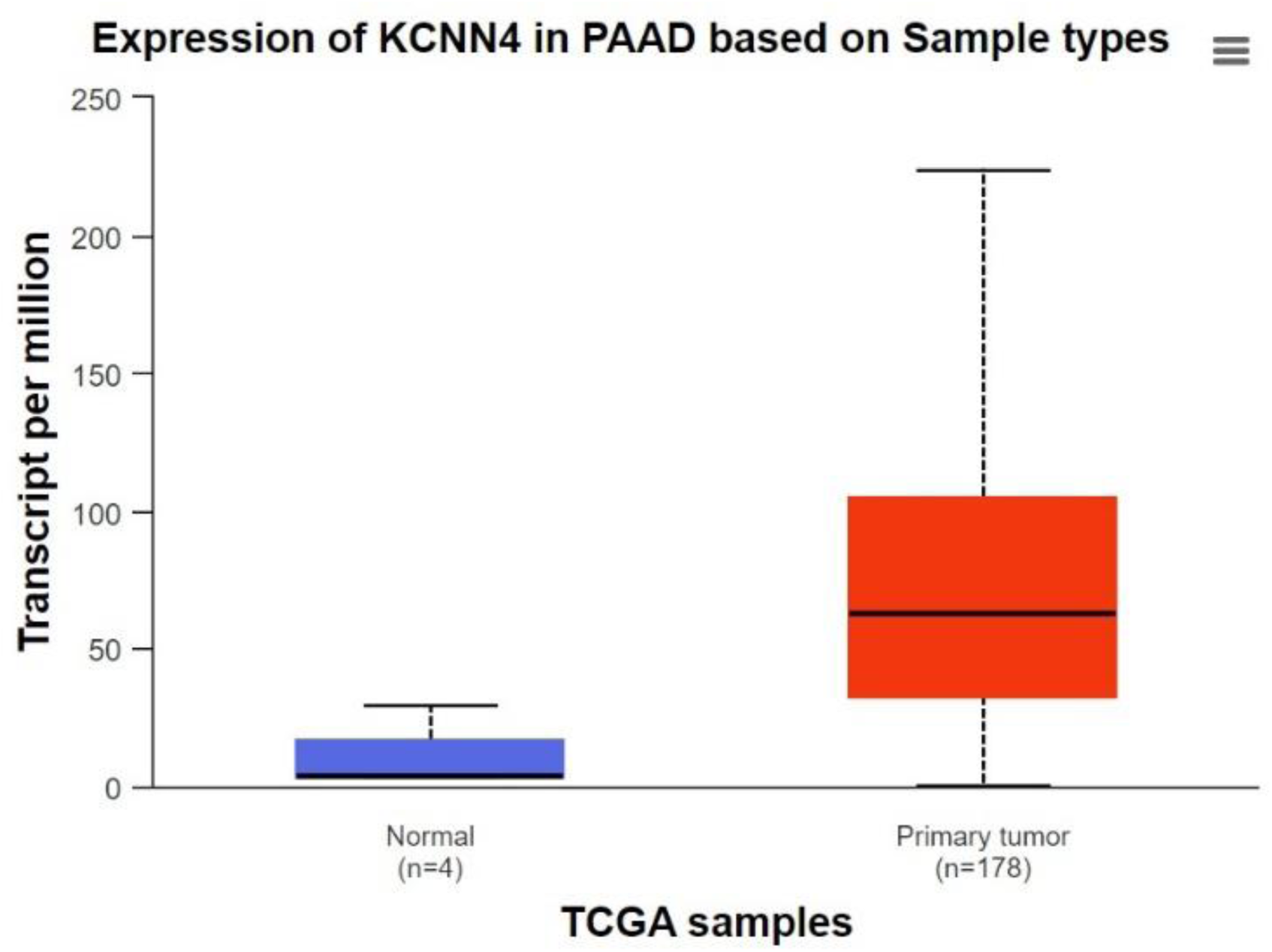
The UALCAN database also indicated that *KCNN4* expressed at a higher level in TCGA PAAD samples compared to nearby normal pancreatic tissue**s.** the analytical report indicated that breast tumor tissues (n=178) had higher levels of KCNN4 expression than normal tissues (n=4).

**Fig 9:**
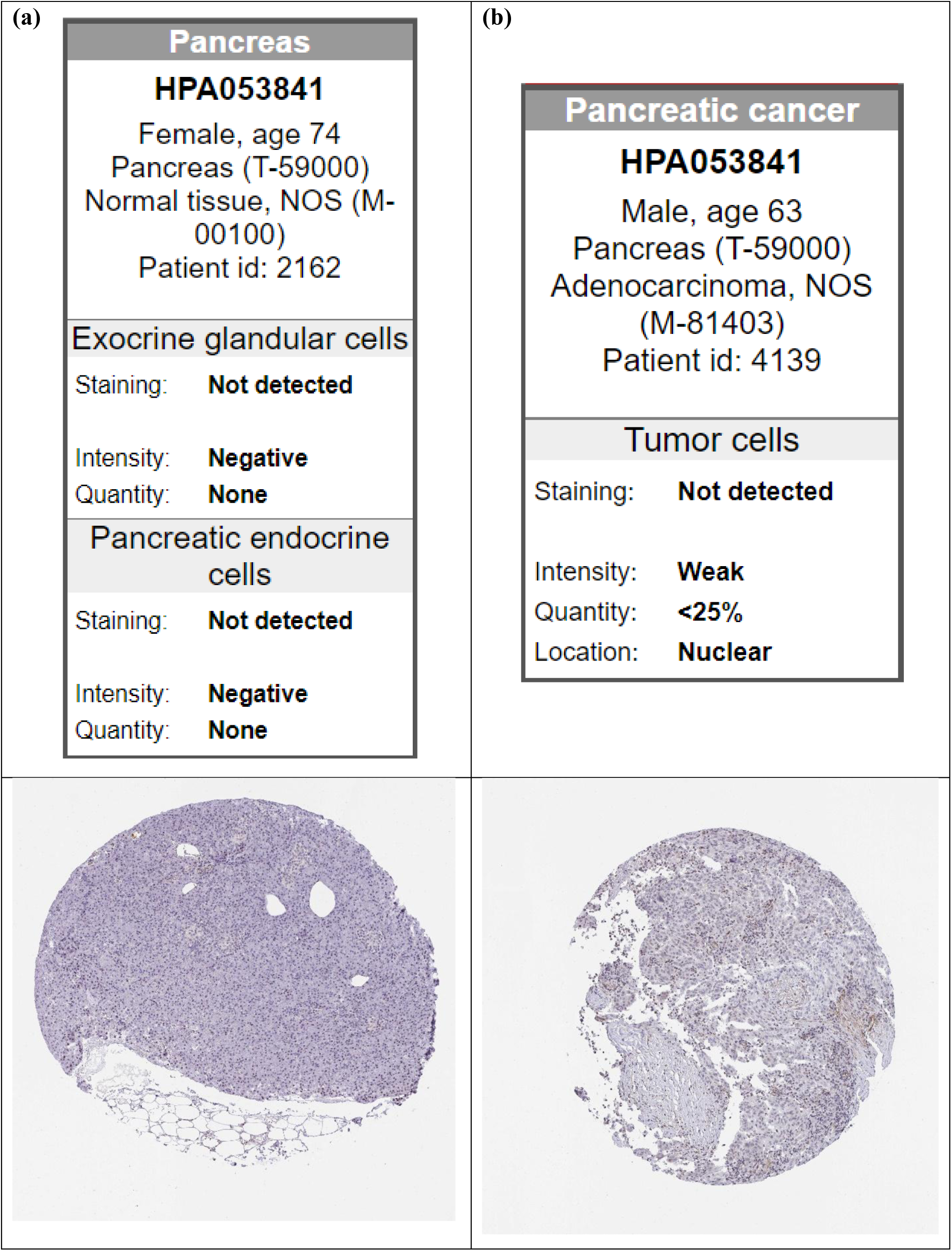
The results of comparative immunohistochemistry also revealed that normal pancreatic tissues had reduced expression of the KCNN4 protein. Moreover, as demonstrated by differing staining intensities from the HPA database, comparative immunohistochemistry observation revealed reduced KCNN4 protein expression in normal breast tissues **(a)** compared to pancreatic cancer tissues **(b)**.

### 3.3 Analysis of the *KCNN4* promoter and DNA-methylation in healthy and diseased pancreatic tissue

Investigation of the promoter methylation of the KCNN4 gene by the UALCAN web server in the TCGA samples suggested that there may be more hypomethylation of the gene found in tumor specimens (n=184) than in normal specimens (n=10) (Figure 10). This finding suggests that *KCNN4* overexpression in pancreatic cancer patients may be related to this, since the continuous activation of the gene is fueled by hypomethylation in the CpG-island of promoters.

**Fig 10:**
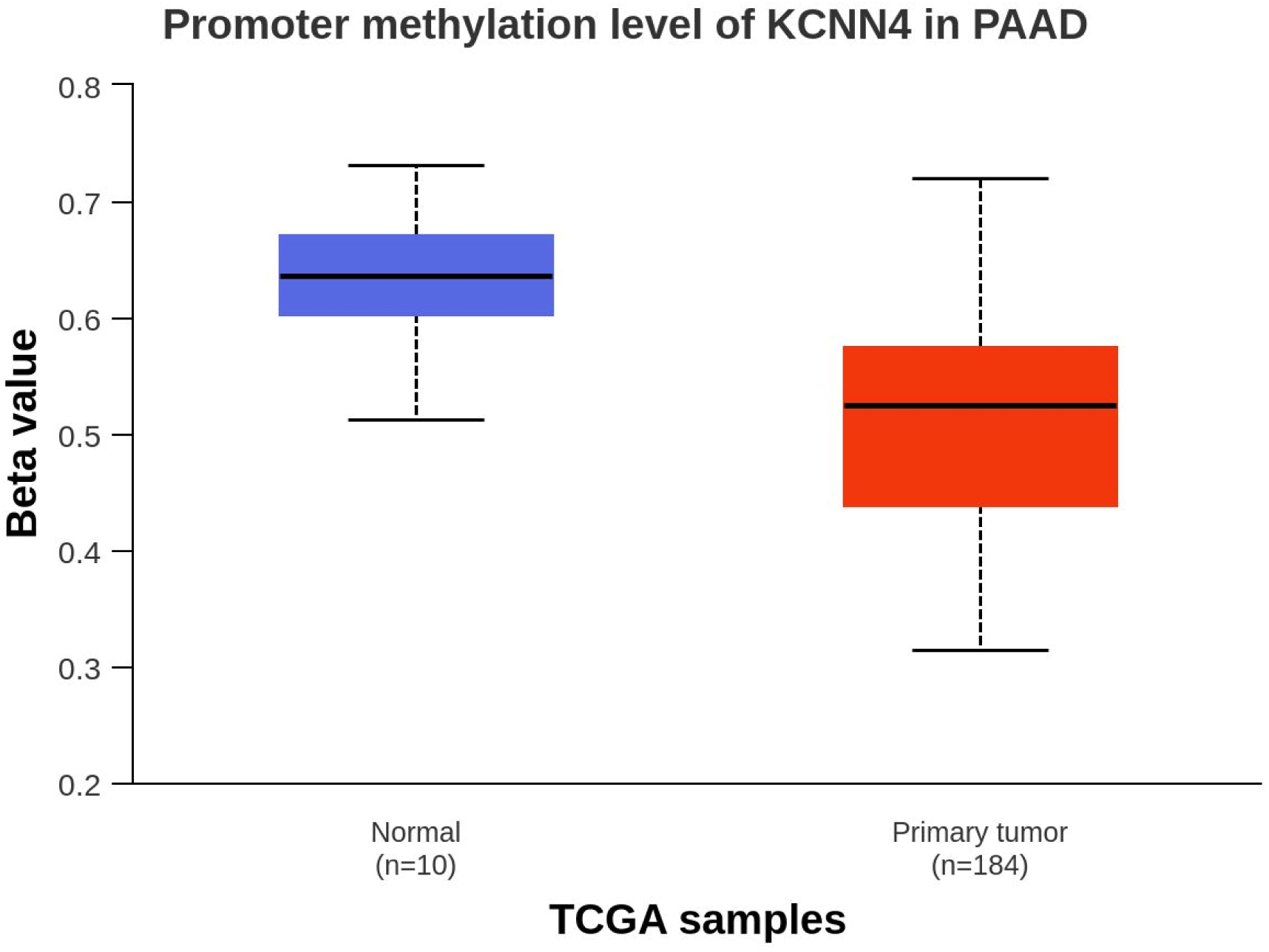
Investigation of the promoter of the *KCNN4* gene and DNA methylation in samples of pancreatic cancer. The UALCAN server revealed the *KCNN4* promoter’s hypomethylation sign.

Web-based tools for clinical and phenotypic annotation of a desired gene are provided by the UCSC Xena platform. The UCSC Xena browser determined the gene expression, copy numbers, and the *KCNN4* gene’s somatic mutations in the GDC TCGA Pancreatic Cancer (PAAD) samples (n=223). Moreover, we also analyzed the neoplasm histologic grade, disease type, and age at initial pathogenesis diagnosis in pancreatic adenocarcinoma samples (Figure 11). From this analysis, we found that the primary tumor was 83%, solid tissue normal was 16.6%, and metastasis was 0.442% among all the samples (n=223).

**Fig 11:**
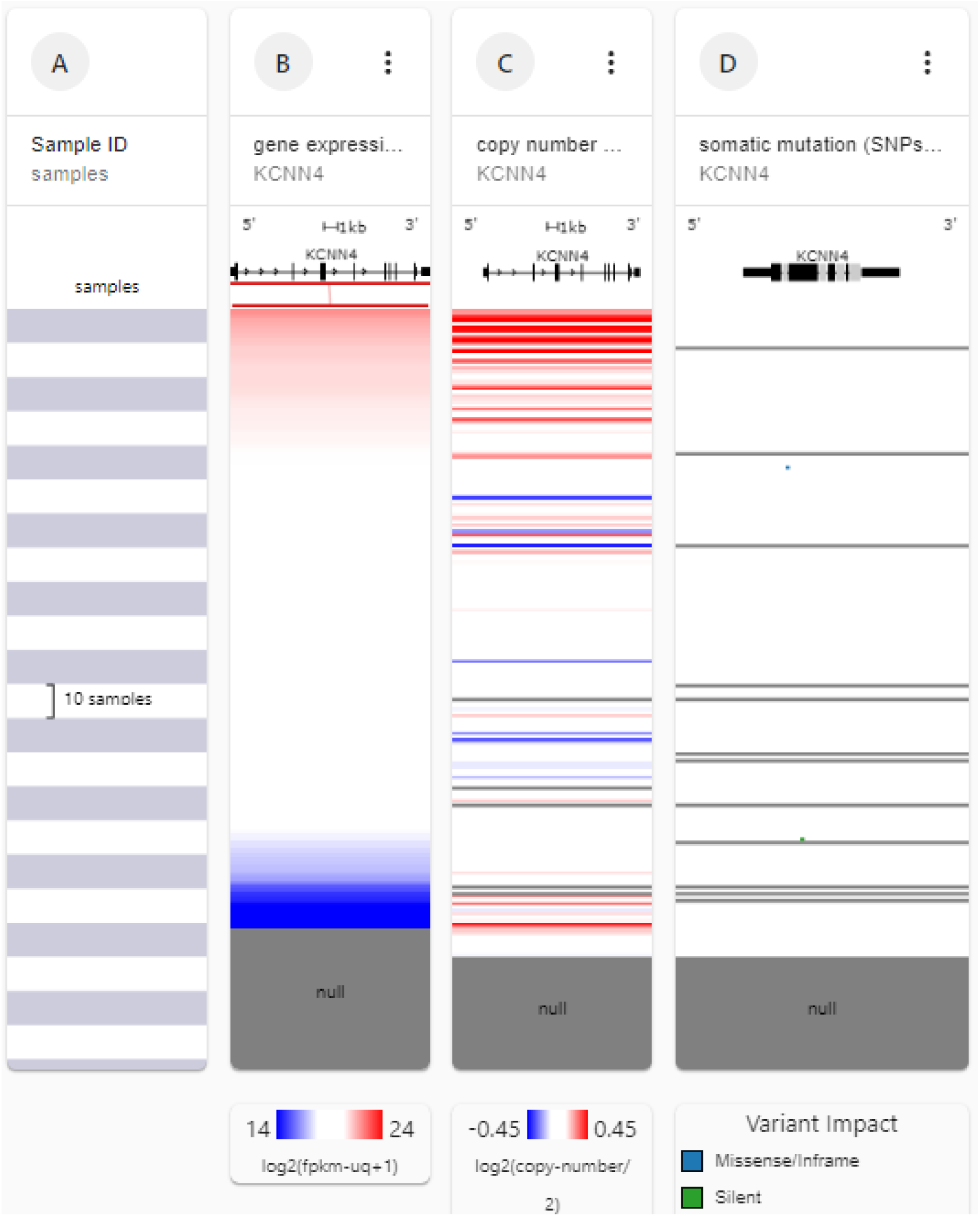
UCSC Xena browser analyzed the gene expression, copy numbers, and the *KCNN4* gene’s somatic mutations in the GDC TCGA Pancreatic Cancer (PAAD) samples.

Among the GDC TCGA Pancreatic Cancer (PAAD) samples the ductal and lobular neoplasms were 81.6%, adenomas and adenocarcinomas 15.2%, cystic, mucinous and serous neoplasm was 2.69%, epithelial neoplasms was 0.448% (Figure 12). A bar chart was formed, wherein the x axis denoted the neoplasm histologic grad and the y axis denoted the age at initial pathologic diagnosis (Figure 13). It was found that neoplasm histologic grade-2 was initially diagnosed at the age of 65 (median). And neoplasm histologic grade-1, grade-3, and grade-4 were initially diagnosed at the age 65, 67, 74 (median) respectively.

**Fig 12:**
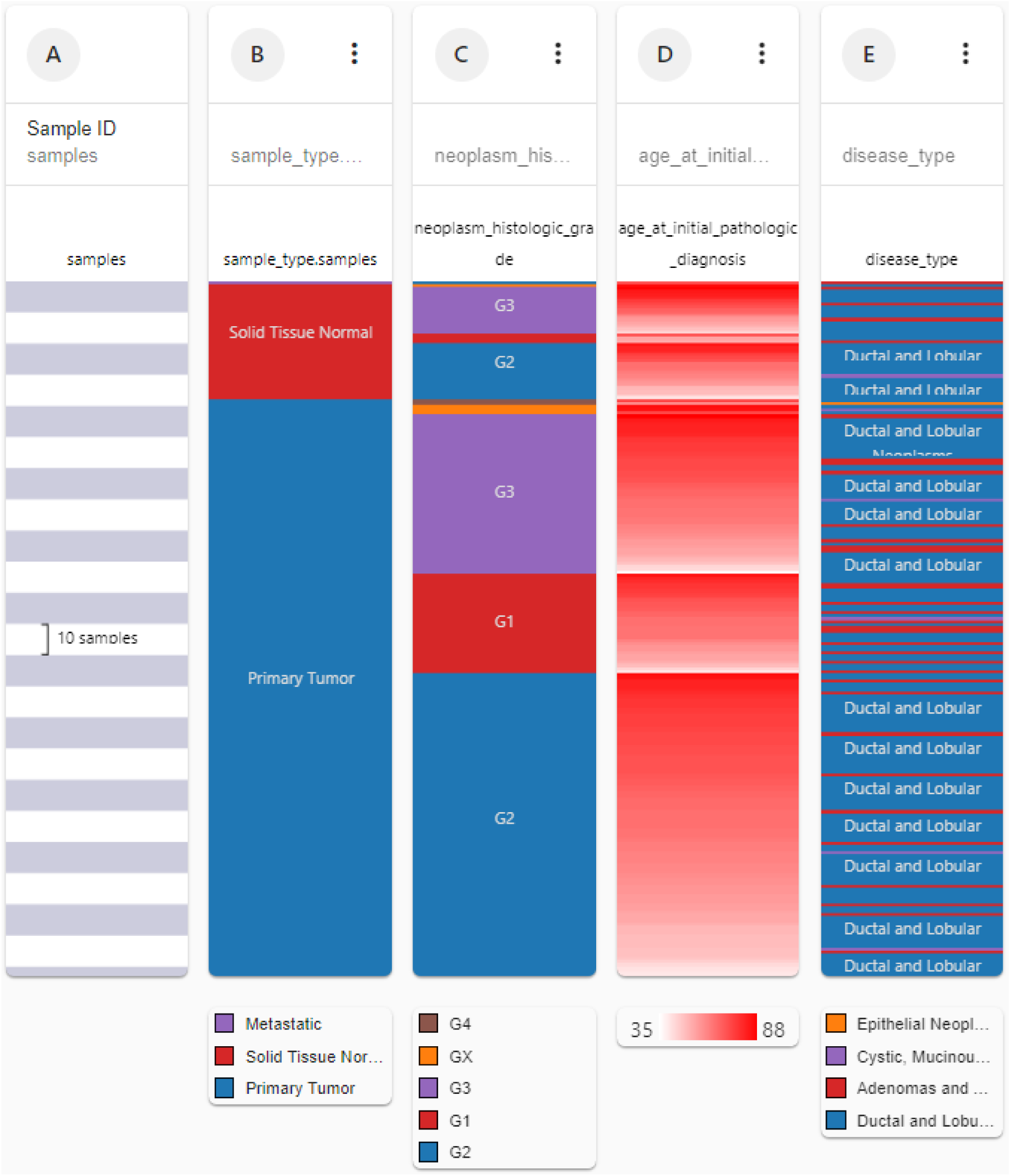
Analysis of the neoplasm histologic grade, disease type, and age at initial pathogenesis diagnosis in PAAD samples by the UCSC Xena browser.

**Fig 13:**
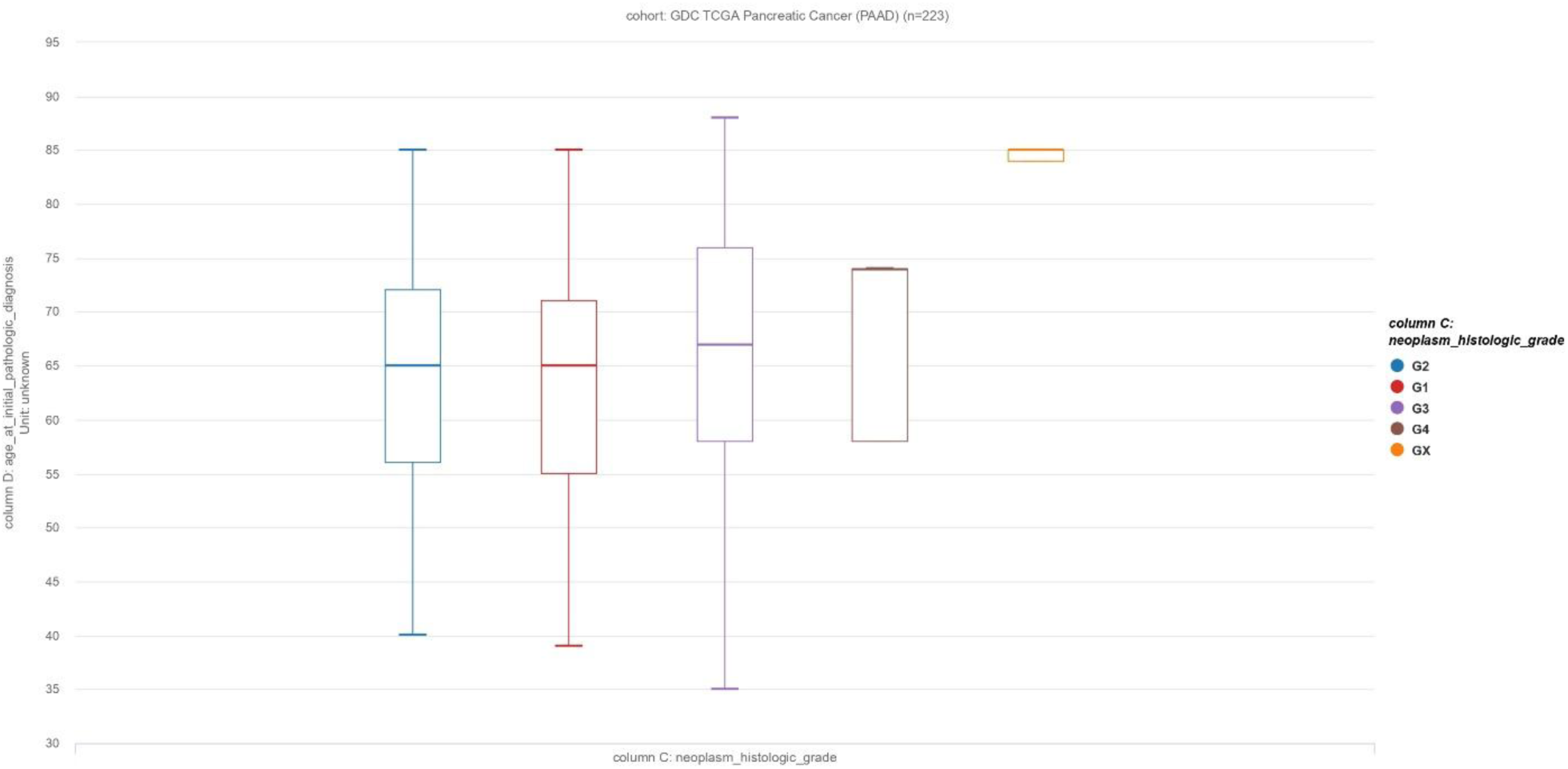
The bar chart represents the relation among histologic grades and initial diagnosis age.

### 3.4 Analyzing the *KCNN4* genes’ copy numbers alteration and mutation frequencies in pancreatic cancer specimens

In pancreatic cancer research, copy numbers alteration and mutations analysis of the KCNN4 gene showed the existence of two somatic mutations with a percent mutation-frequency of 0.2 in all of the pancreatic cancer samples that were chosen for assessment (Tab 1). Of the various kinds of mutations found in the pancreatic cancer study samples, such as amplification, deep-deletion, and nonsense, the missense mutations were the most common form (2) (Tab 1). On chromosome 19, every mutation fell between 44278467 bp (start) and 44280710 bp (end), in charge of encoding the KCNN4 protein, which has 427 amino acids (Fig 14 and Table 1). The pancreatic cancer sample (UTSW) showed the highest alteration frequency; in 14.68% of the 109 cases, the gene was altered. The gene was changed in 2.72% of the 184 cases in the pancreatic adenocarcinoma (TCGA) sample; of these, 0.54% (1 case) had a mutation and 2.17% (4 cases) had an amplification. Finally, the gene changed in 1.02% of the 98 cases of pancreatic neuroendocrine tumors. When combined, these genetic differences appear to be important factors in the prognosis of pancreatic adenocarcinoma (PAAD) (Fig 15).

**Table 1:**
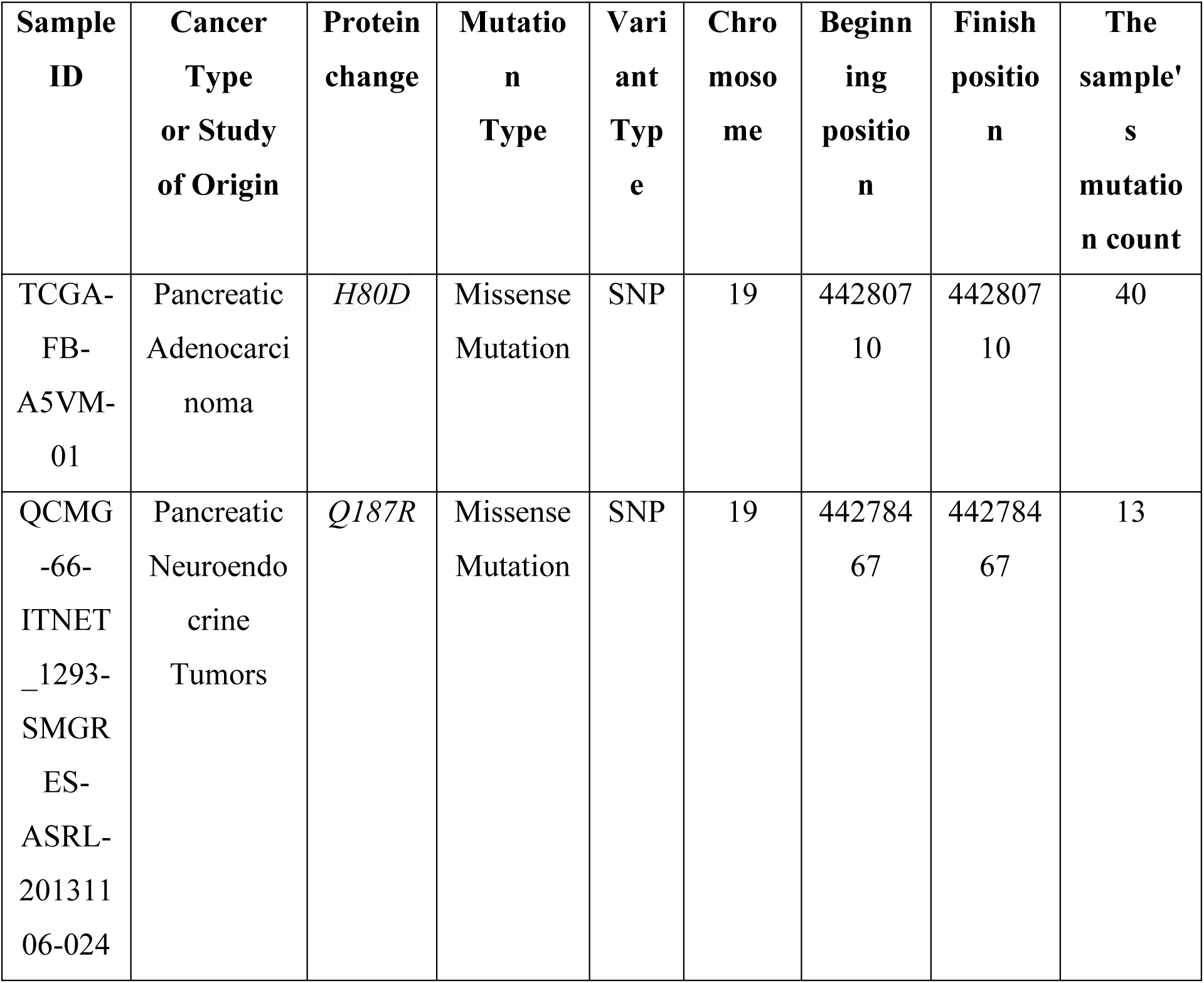
Summary of the KCNN4 gene mutations and copy numbers alteration analysis in pancreatic cancer samples.

| **Sample ID** | **Cancer Type or Study of Origin** | **Protein change** | **Mutation Type** | **Variant Type** | **Chromosome** | **Beginning position** | **Finish position** | **The sample's mutation count** |
| --- | --- | --- | --- | --- | --- | --- | --- | --- |
| TCGA-FB-A5VM-01 | Pancreatic Adenocarcinoma | *H80D* | Missense Mutation | SNP | 19 | 44280710 | 44280710 | 40 |
| QCMG-66-ITNET_1293-SMGR-ES-ASRL-20131106-024 | Pancreatic Neuroendocrine Tumors | *Q187R* | Missense Mutation | SNP | 19 | 44278467 | 44278467 | 13 |

**Fig 14:**
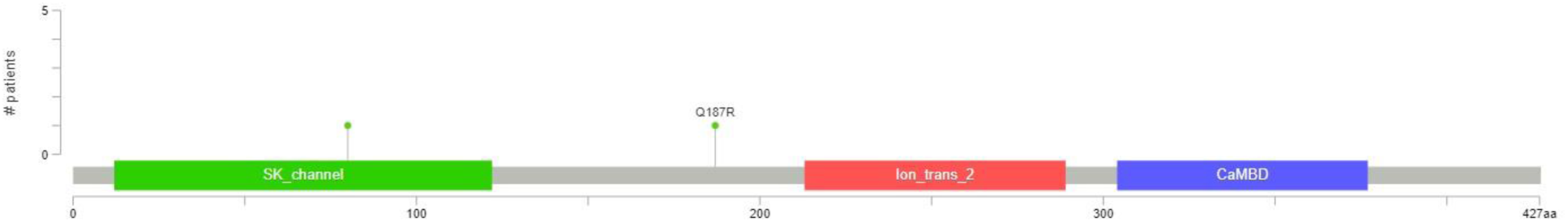
The frequency of copy number alteration and *KCNN4* mutation in various pancreatic cancer research. A lollipop diagram showing the common mutations found in pancreatic cancer samples’ *KCNN4* gene.

**Fig 15:**
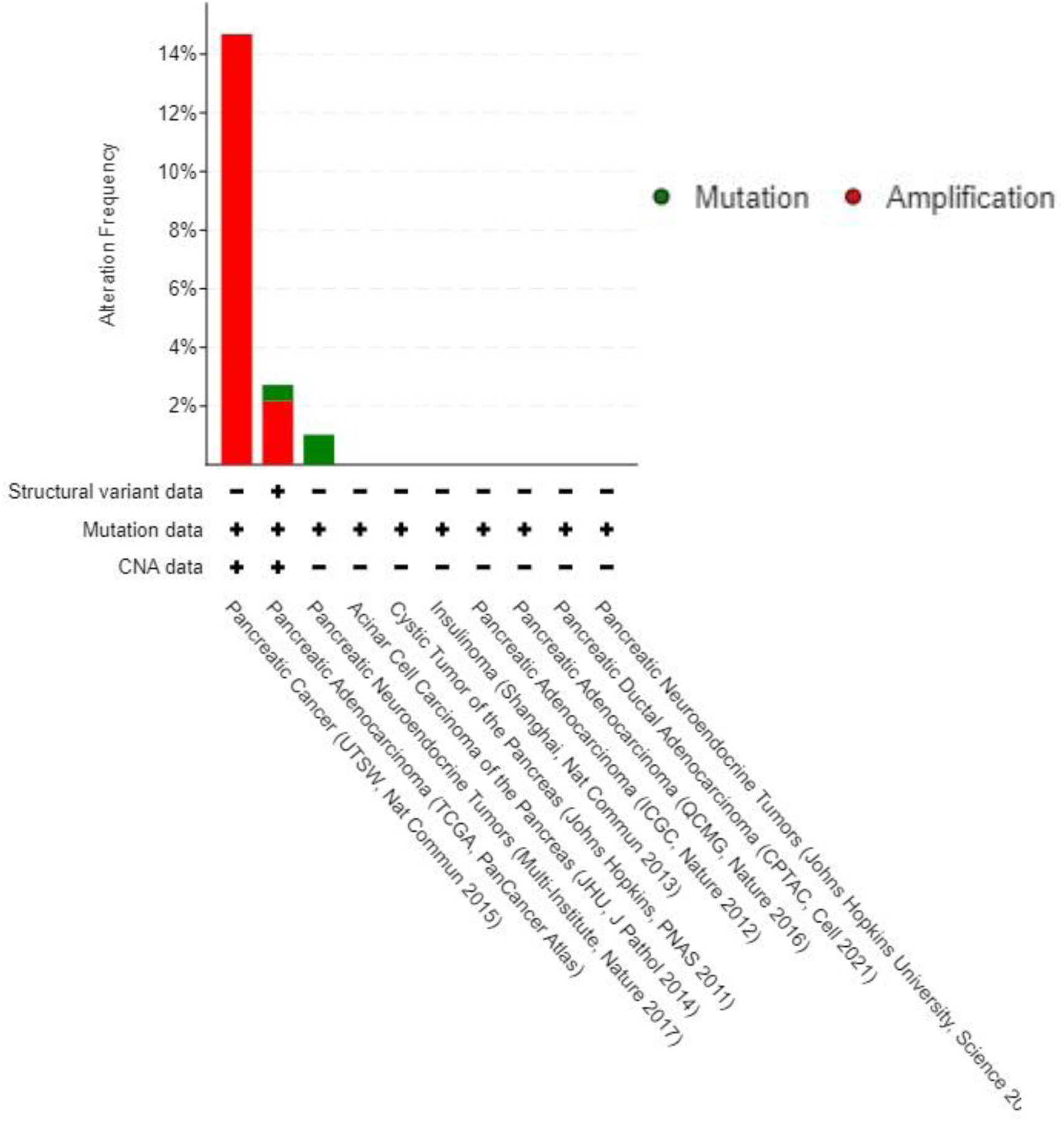
Bar diagram showing the various patterns of genetic variations in the *KCNN4* gene that has been discovered in a number of studies on pancreatic cancer.

### 3.5 Relationship between pancreatic cancer patients’ clinical characteristics and *KCNN4* gene expression

With the help of the UALCAN server, the association between various clinical characteristics of patients with pancreatic cancer, such as age, sex, race, and *KCNN4* expression, was assessed using the TCGA sample (Table 2 and Figure 16). The most noticeable overexpression was observed in tumor grade 2 (p =7.005700E-02) in relation to other tumor-grades found in the TCGA specimens (Figure 16e). Significant and varying levels of KCNN4 expression were also observed in all the chosen clinical characteristics of patients with pancreatic cancer. Remarkably, across the age ranges, it was discovered that middle-aged patients (41–60 years) expressed more *KCNN4* than younger (21-40) and old-age patients (61-100 years) (Figure 16d). It was also discovered that expression increased markedly as the cancer stage progressed, peaking at stage-4 of pancreatic adenocarcinoma (PAAD) (Figure 16a). Men showed higher levels of *KCNN4* expression than women in terms of gender-specific stratification. (Figure 16c). Furthermore, compared to patients of other racial categories, Asians displayed a higher degree of expression. Furthermore, compared to TP53 non mutant pancreatic cancer patients, TP53 mutant pancreatic cancer patients were found to express higher levels of *KCNN4*. Taken together, these results indicate that variations in *KCNN4* expression patterns possibly important in how the cancer prognosis manifests and, as a result, positively correlate with various clinical characteristics of patients with pancreatic cancer.

**Table 2:**
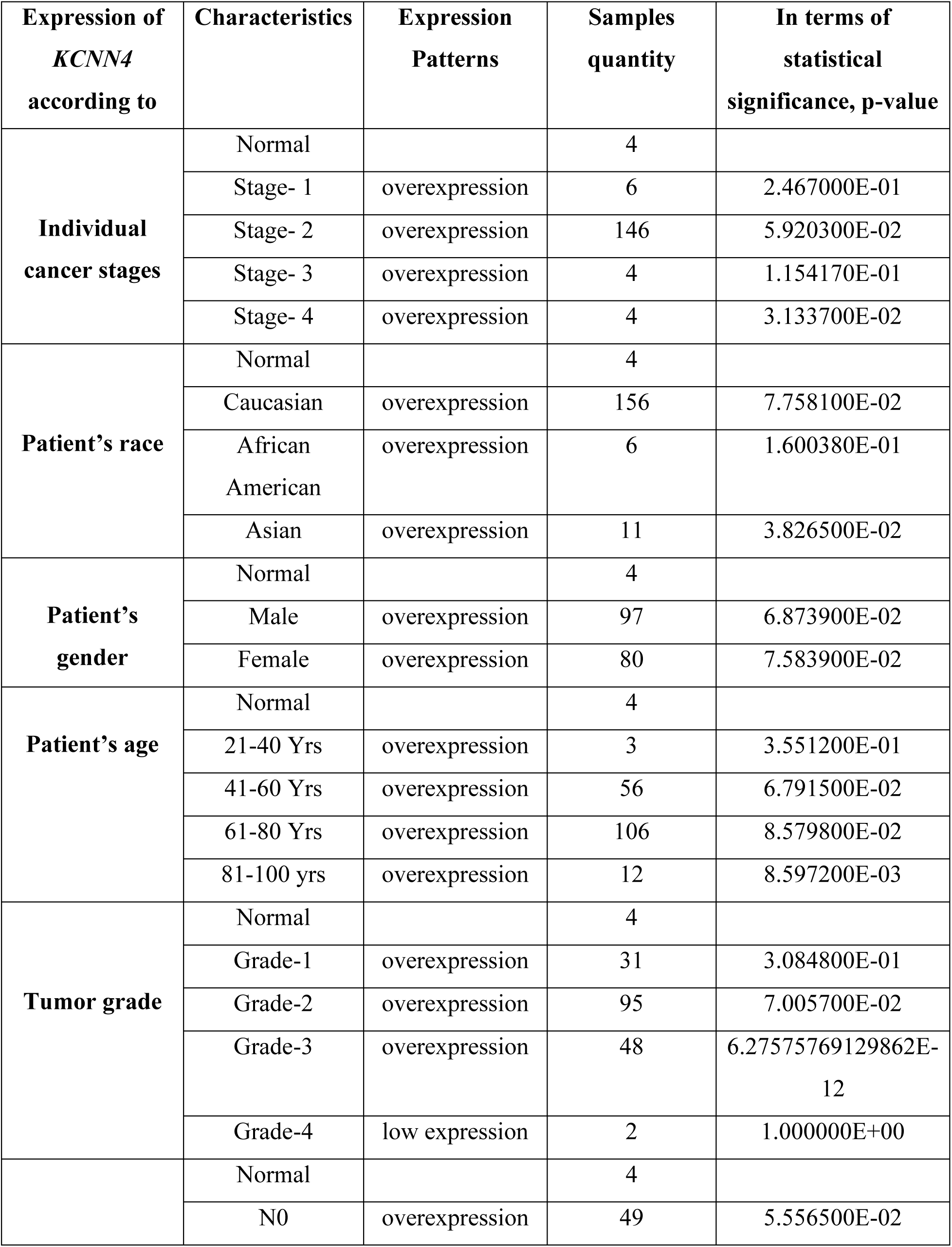
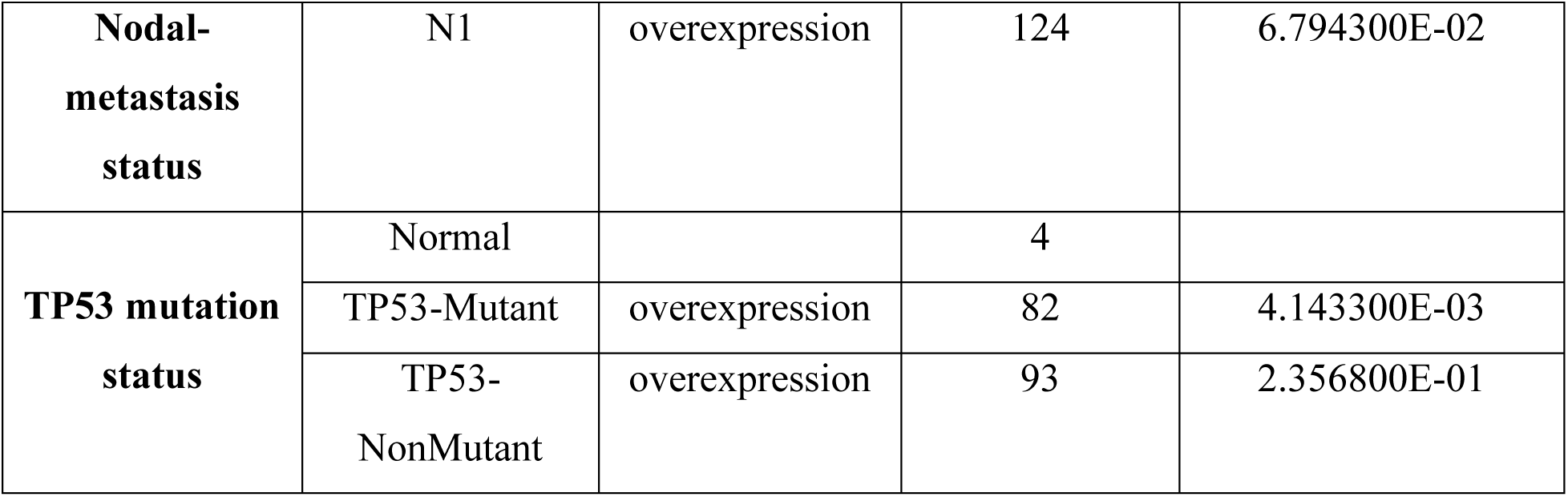
An analysis overview of copy-number alterations and *KCNN4* mutations in PAAD samples.

**Fig 16:**
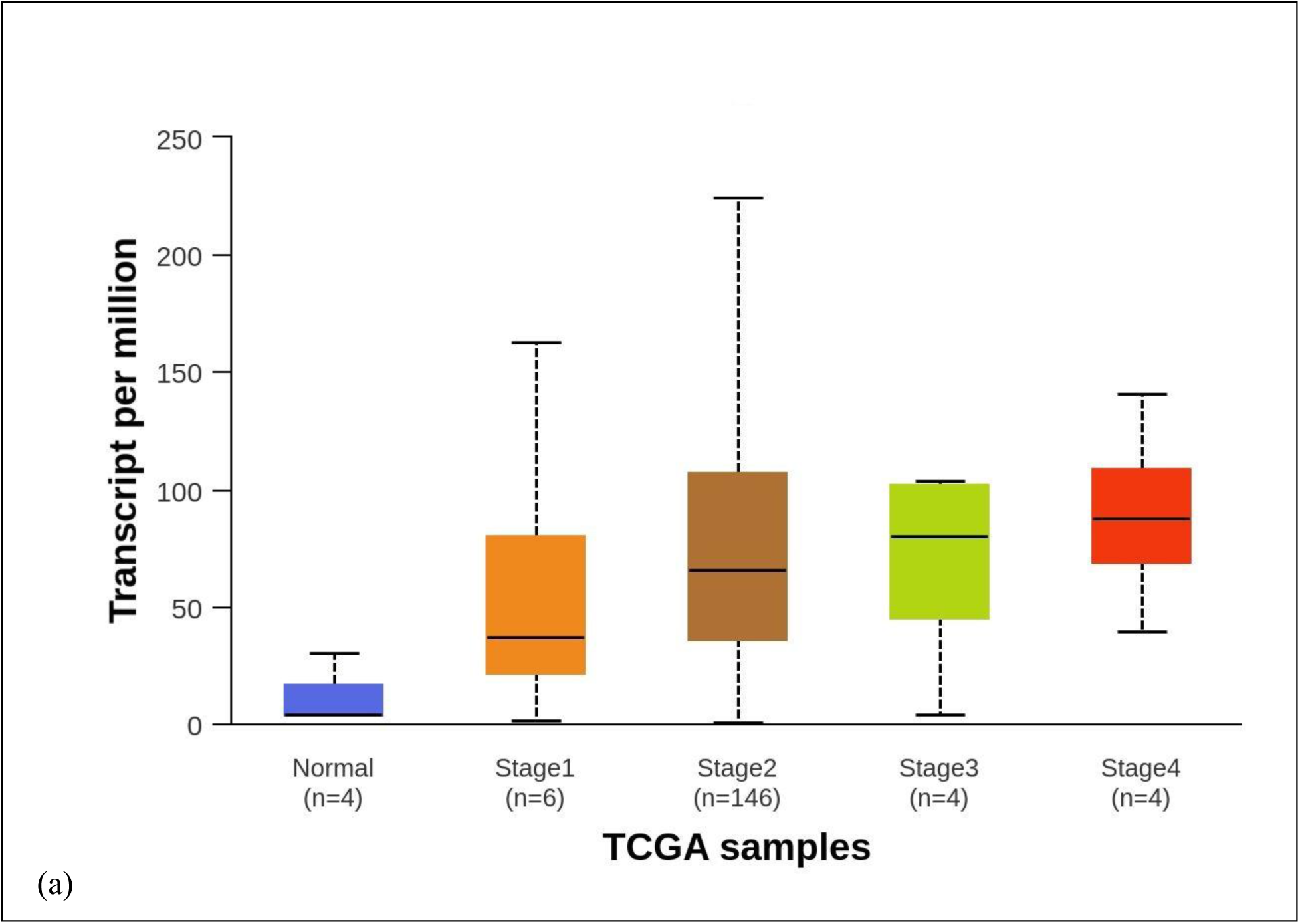
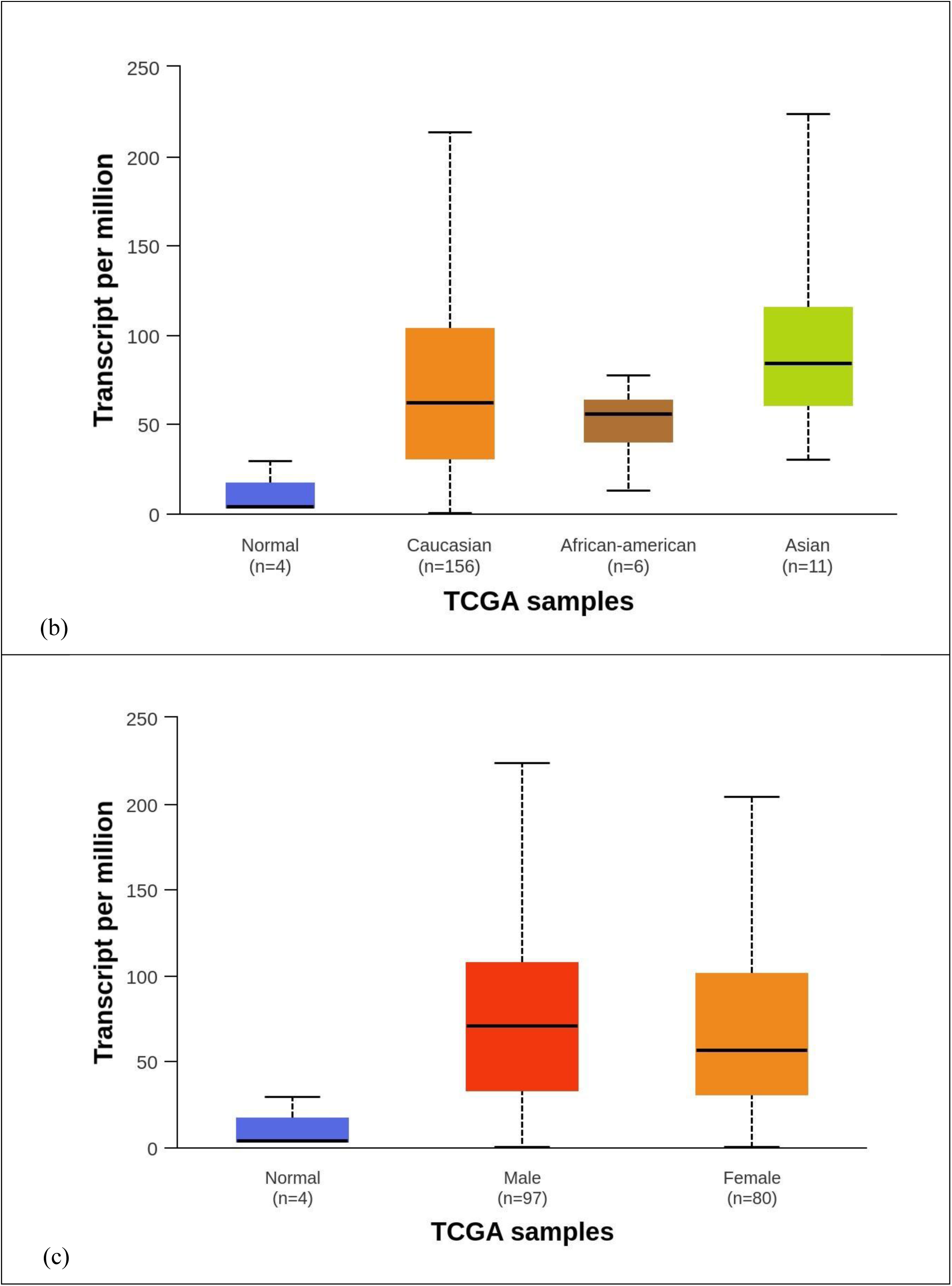
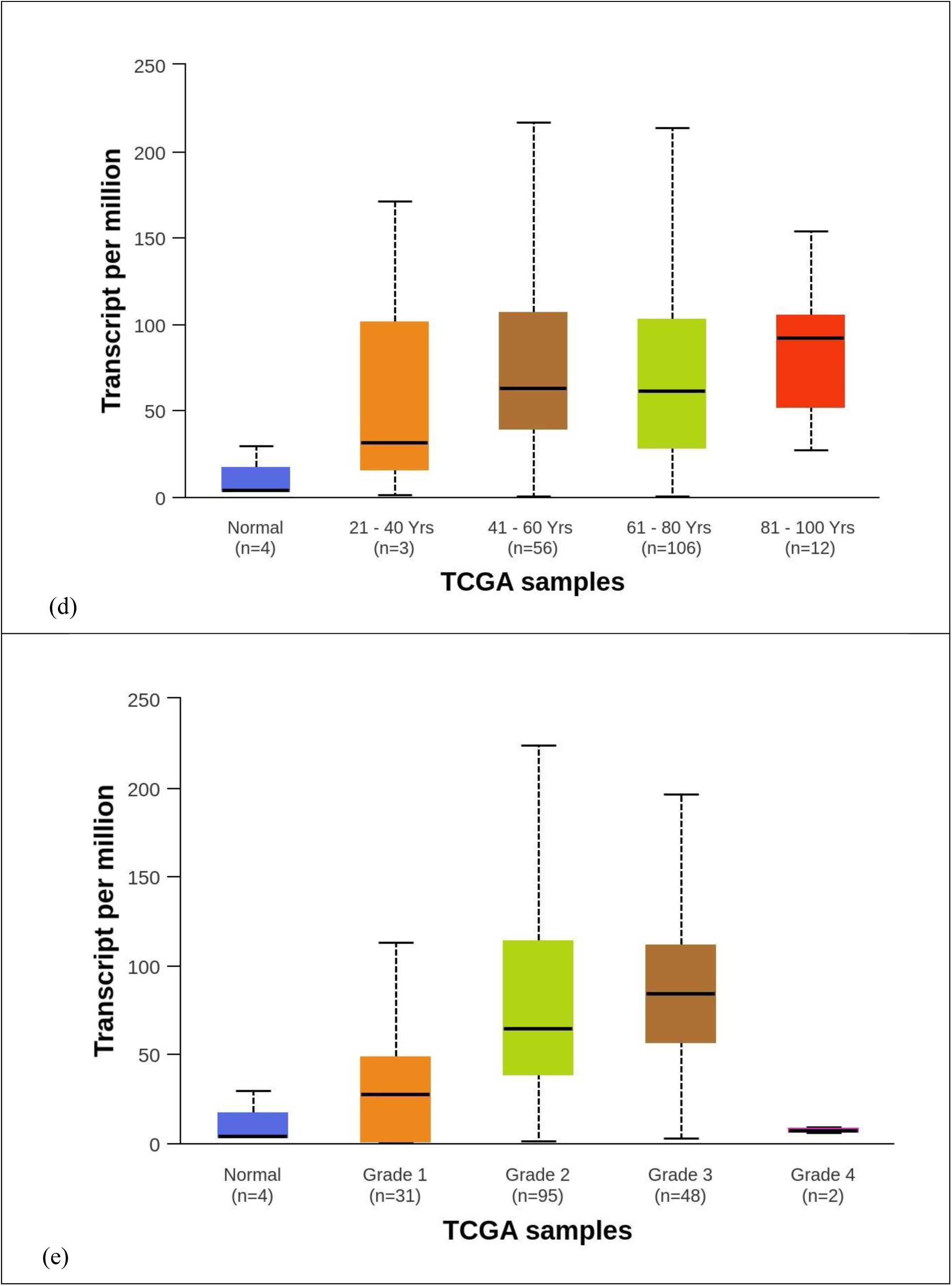
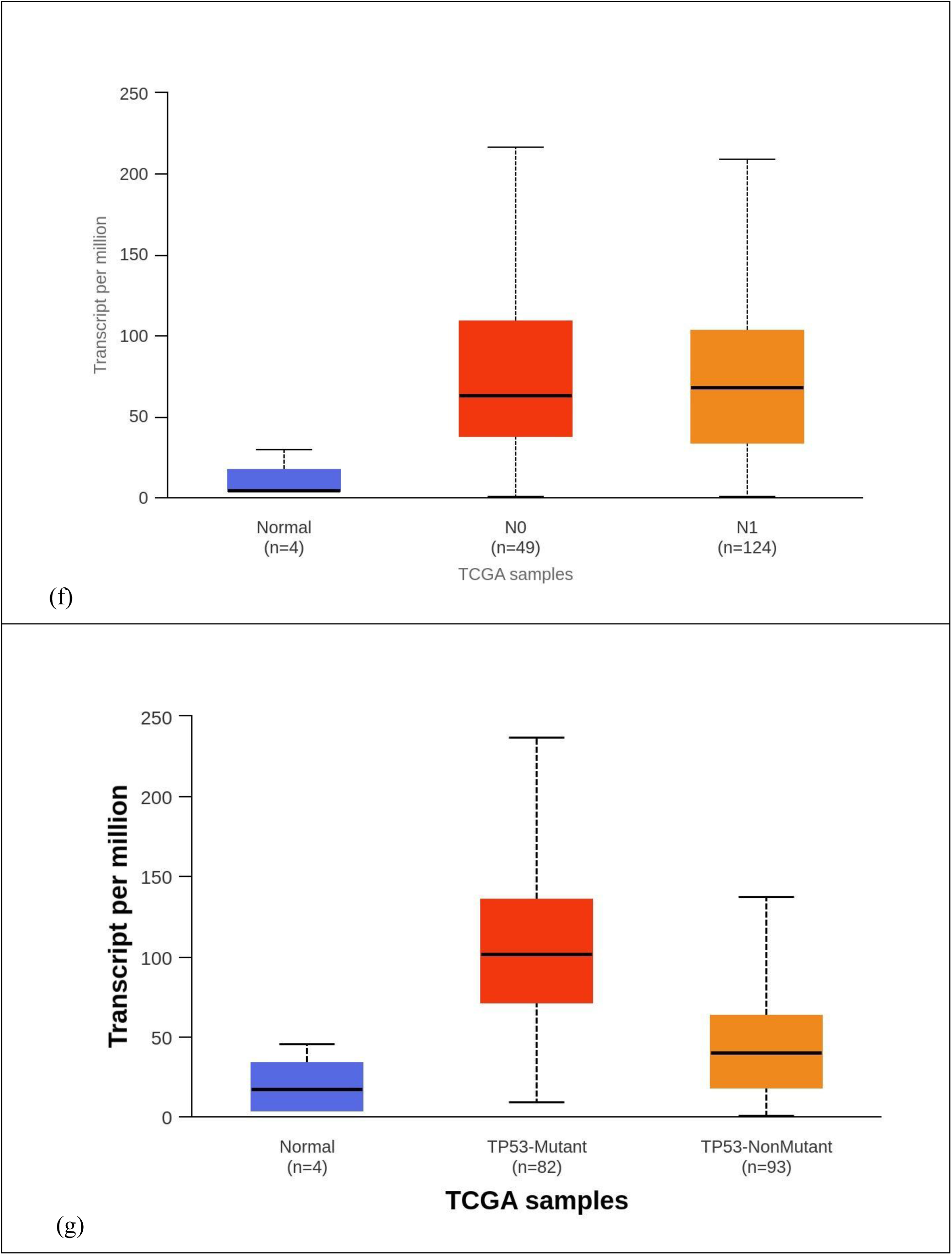
Boxplot showing the relationship between multiple clinical features of pancreatic adenocarcinoma (PAAD) patients and *KCNN4* gene expression. The following factors were analyzed: (a) stage of cancer; (b) race of patients; (c) gender of patients; (d) age of patients; (e) grade of tumor; (f) nodal-metastasis; and (g) status of TP53 mutation. KCNN4 expression was found to be elevated in all of the needed clinical parameters.

### 3.6. The survival trend of patients with pancreatic cancer in the response to *KCNN4* **Expression**

Variations in patient survival rates with pancreatic cancer following the expression of *KCNN4* are shown by Kaplan-Meier (KM) plots in Figure 17. According to the analysis, there is a negative correlation between the higher expression of KCNN4 and pancreatic cancer patients’ overall-survival (OS), utilizing a hazard ratio (HR) of 1.19 (p=0.0165) (Figure 17a). Similar types of findings regarding disease-free survival (DFS) were also made (HR=0.57; p=<4.37*10^-05^) (Figure 17b). The results showed that disease-free survival (DFS) patients had a lower survival rate than overall-survival OS patients, with a median life expectancy of 60–120 months.

When considered collectively, these collected data indicate that increased *KCNN4* expression may be linked to a decreased life expectancy in pancreatic cancer patients and may be a predictor of their survival status.

**Fig 17:**
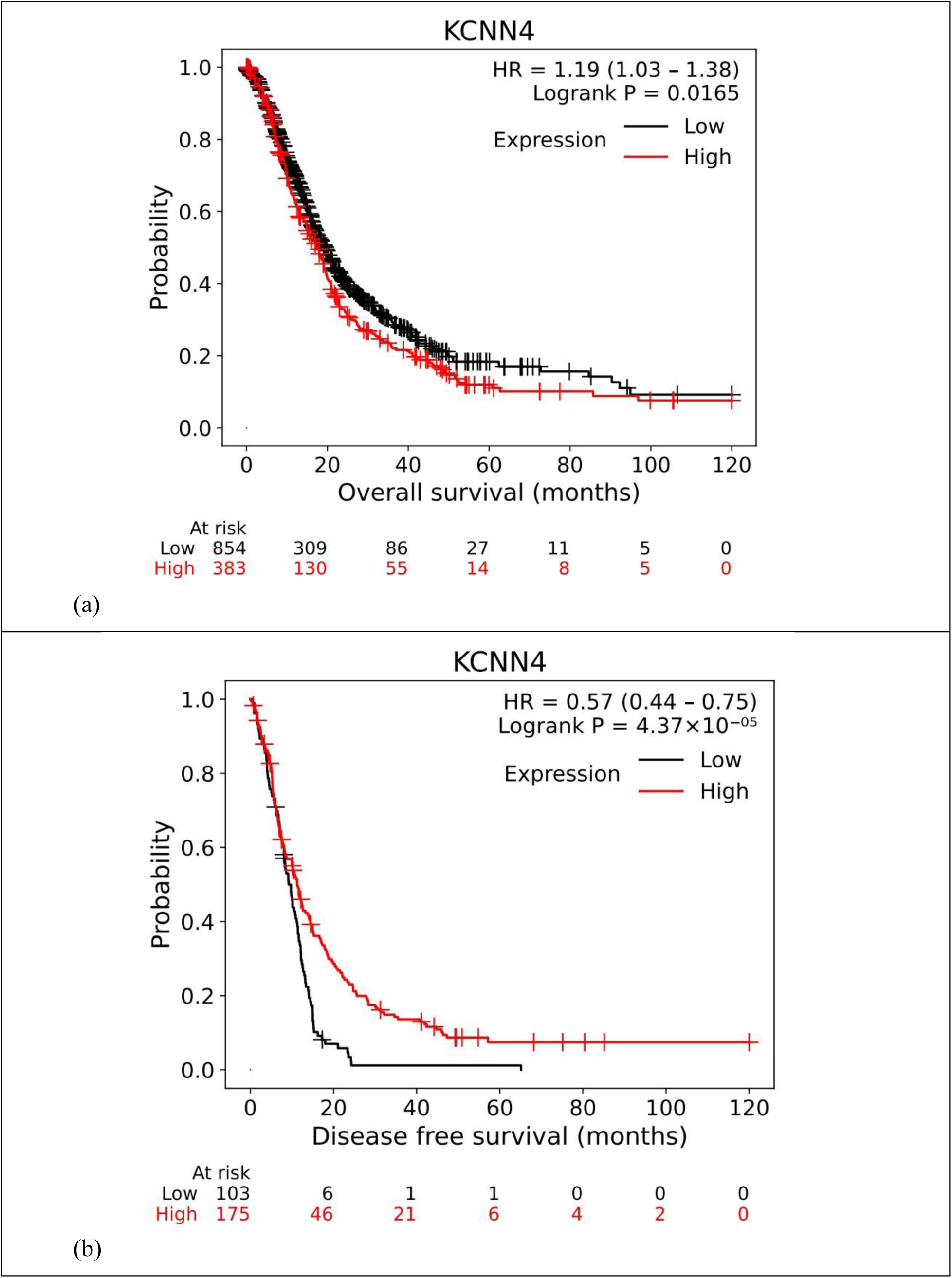
Kaplan-Meier plots illustrating the predicted survival trend of patients with PAAD versus KCNN4 expression. **(a)** Overall survival; **(b)** Disease free survival. There was a negative correlation between the expression of *KCNN4* and the different survival rates of PAAD patients.

### 3.7 Analysis of genes functionally similar to *KCNN4* in pancreatic adenocarcinoma

Understanding the *KCNN4*’s fundamental regulatory mechanism in pancreatic adenocarcinoma cells, the PAAD sample in the GEPIA2 server was used to carry out similar gene detection in order to find functionally related genes. *PPP1R13L* (protein phosphatase 1 regulatory subunit 13-like) was discovered as the most likely gene that was similar in various pancreatic adenocarcinoma tissues with a Pearson Correlation Coefficient (PCC) of 0.81 (Table 3).

**Table 3:**
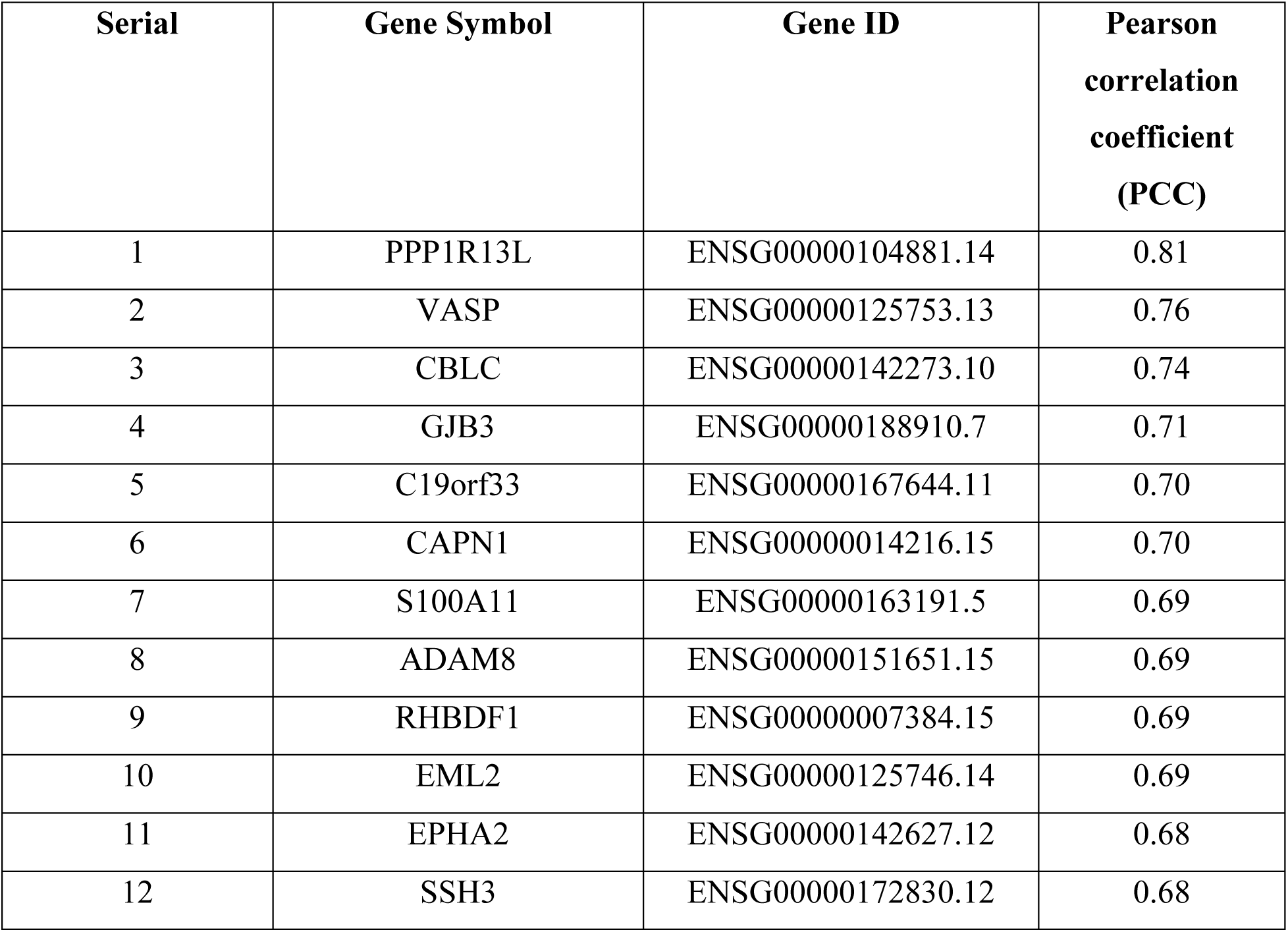
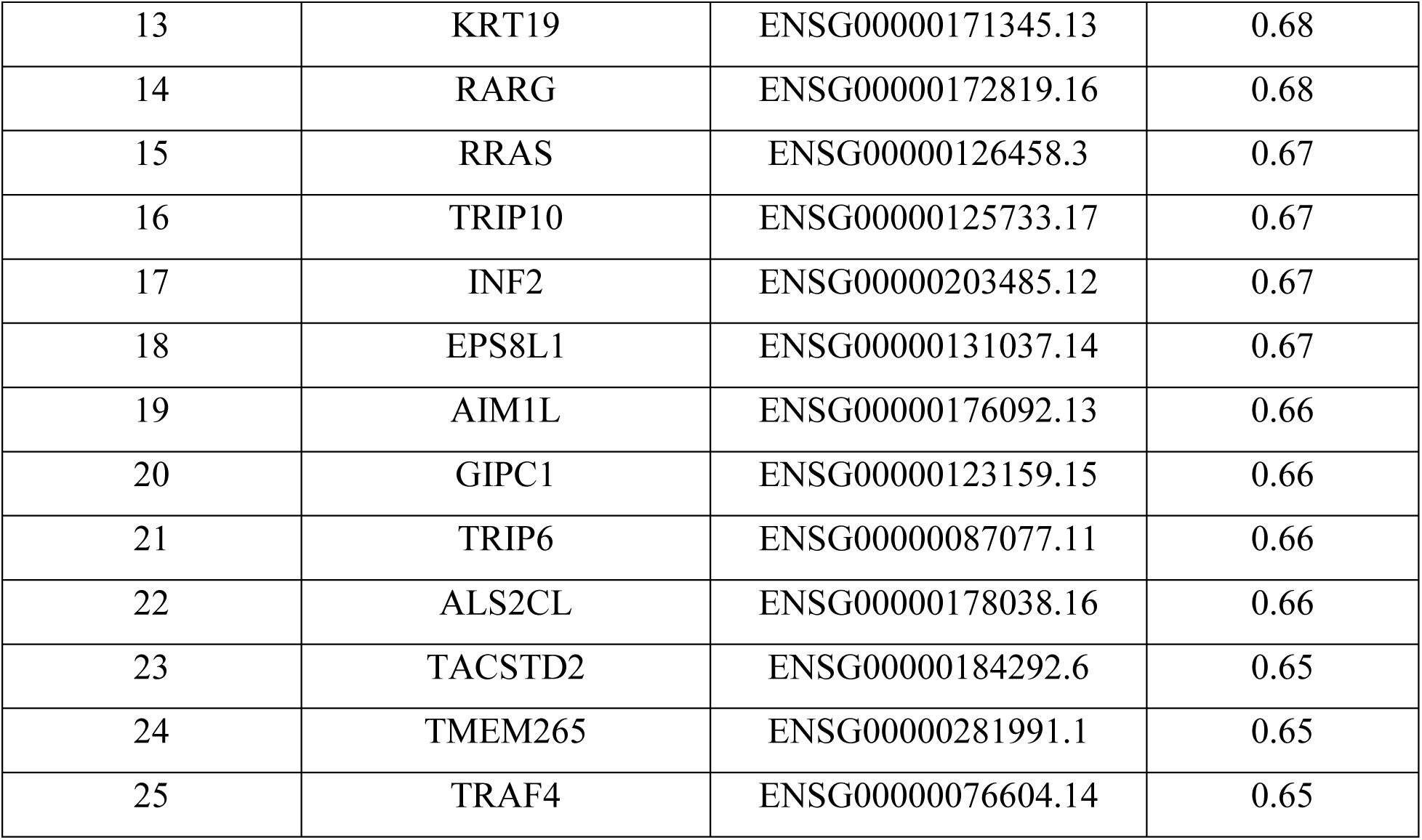
Similar genes with *KCNN4* in PAAD tumor sample from the GEPIA2 server database and ranked them based on their Pearson correlation coefficient (PCC).

Another comparable finding was also made utilizing the GEPIA2 co-expression analysis, which indicated that the gene (*PPP1R13L*) had a 0.81 (p=0) correlation coefficient and was positively co-expressed in PAAD tissues (Figure 18a). Among the other functionally similar genes, *VASP*, *CBLC*, and *GJB3* additionally revealed that the level of *KCNN4* gene expression in pancreatic adenocarcinoma tissues correlated positively (PCC:> 0.70) (Figure 18b, 18c, 18d).

**Fig 18:**
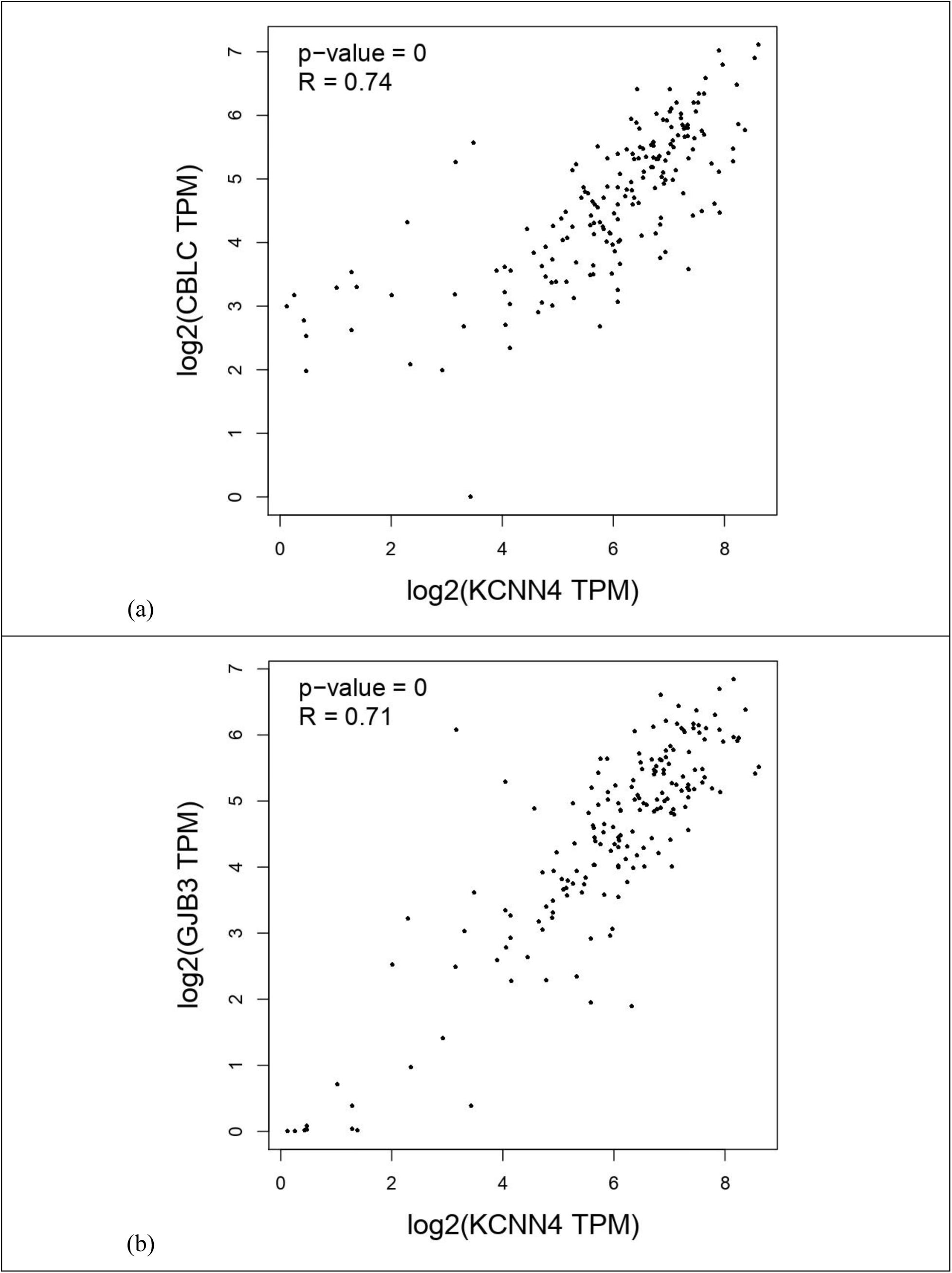
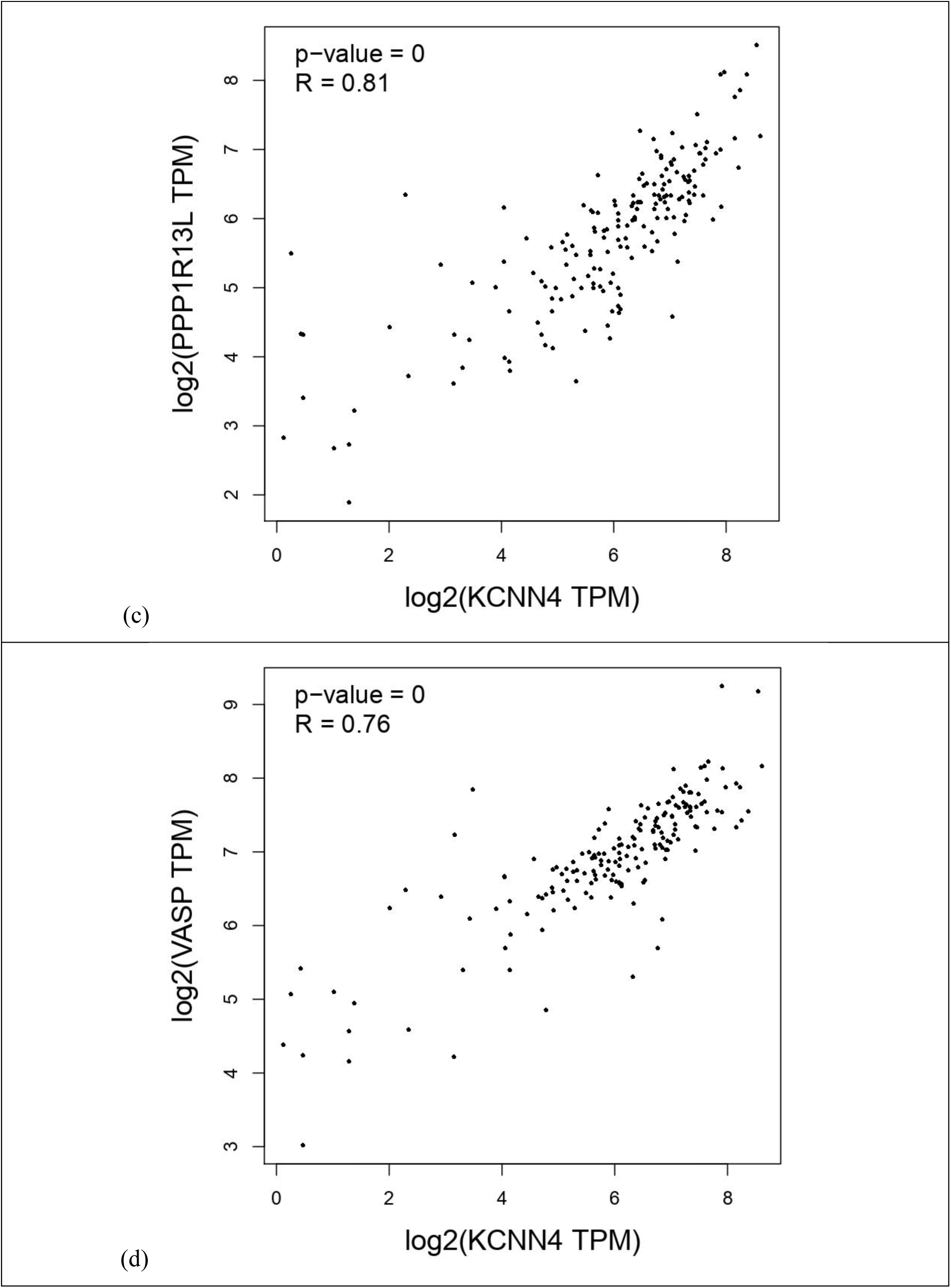
Analysis of co-expression of *KCNN4* and *PPP1R13L* genes using the GEPIA2 database (Fig 18a). Analysis of co-expression of *KCNN4* and *VASP* genes using the GEPIA2 database (Fig 18b). Analysis of co-expression of *KCNN4* and *CBLC* genes using the GEPIA2 database (Fig 18c). Analysis of co-expression of *KCNN4* and *GJB3* genes using the GEPIA2 database (Fig 18d)

### 3.8 Analysis of *KCNN4* co-expressed gene functional enrichment in PAAD tissues

Enrichr web server was used to understand the gene ontologies (GO), which are biological processes, molecular functions, cellular components, and pathways associated with the co-expressed *KCNN4* genes in the pancreatic cancer specimens from the prior step. Analysis of the GO discovered that the majority of the genes have a role in controlling positive regulations of cell differentiation, regulation of ruffle assembly, negative regulation of protein polymerization, regulation of epidermal growth factor receptor signaling pathway, negative regulation of supramolecular fiber organization, cadherin binding, actin binding, and SH3 domain binding (Figures 19a and 19c). Furthermore, the majority of the co-expressed genes of the *KCNN4* in PAAD tissues were found to be primarily involved in focal adhesion, cell-cell junction, cell-substrate junction, and connexin complex, according to cellular component analysis (Figure 19b).

Several pathway analyses, including the Reactome, KEGG, and BioPlanet pathways, were also conducted using the co-expressed genes. The co-expressed genes’ pathway analysis showed that they are primarily in charge of maintaining the TGF-beta regulation of extracellular matrix, integrin signaling pathway, mitochondrial pathway of apoptosis, axon guidance, Rap1 signaling pathway. Cellular senescence, Ras signaling pathway, developmental biology R-HSA-1266738, and so on (Figure 19d, 19e, and 19f).

**Fig 19:**
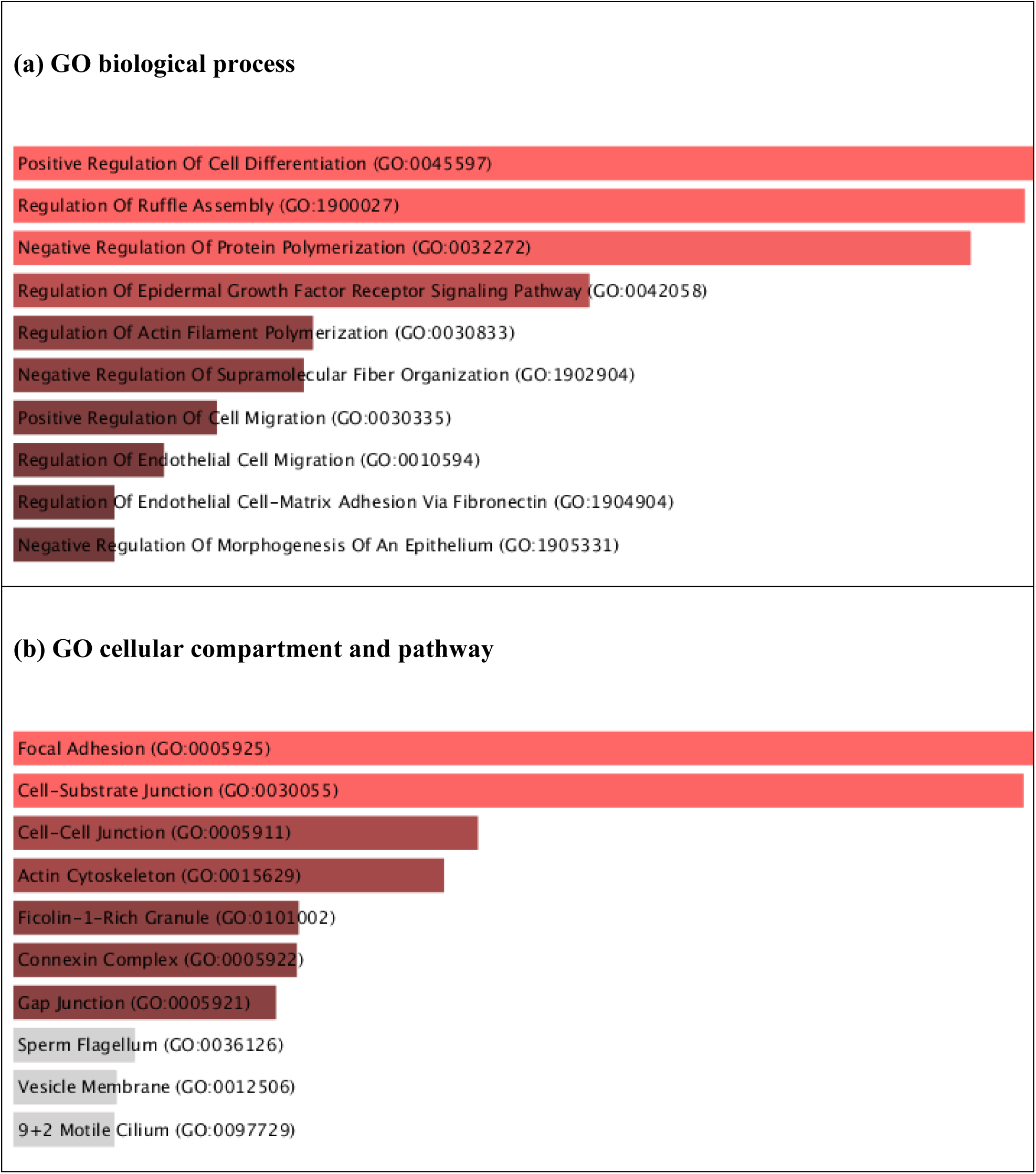
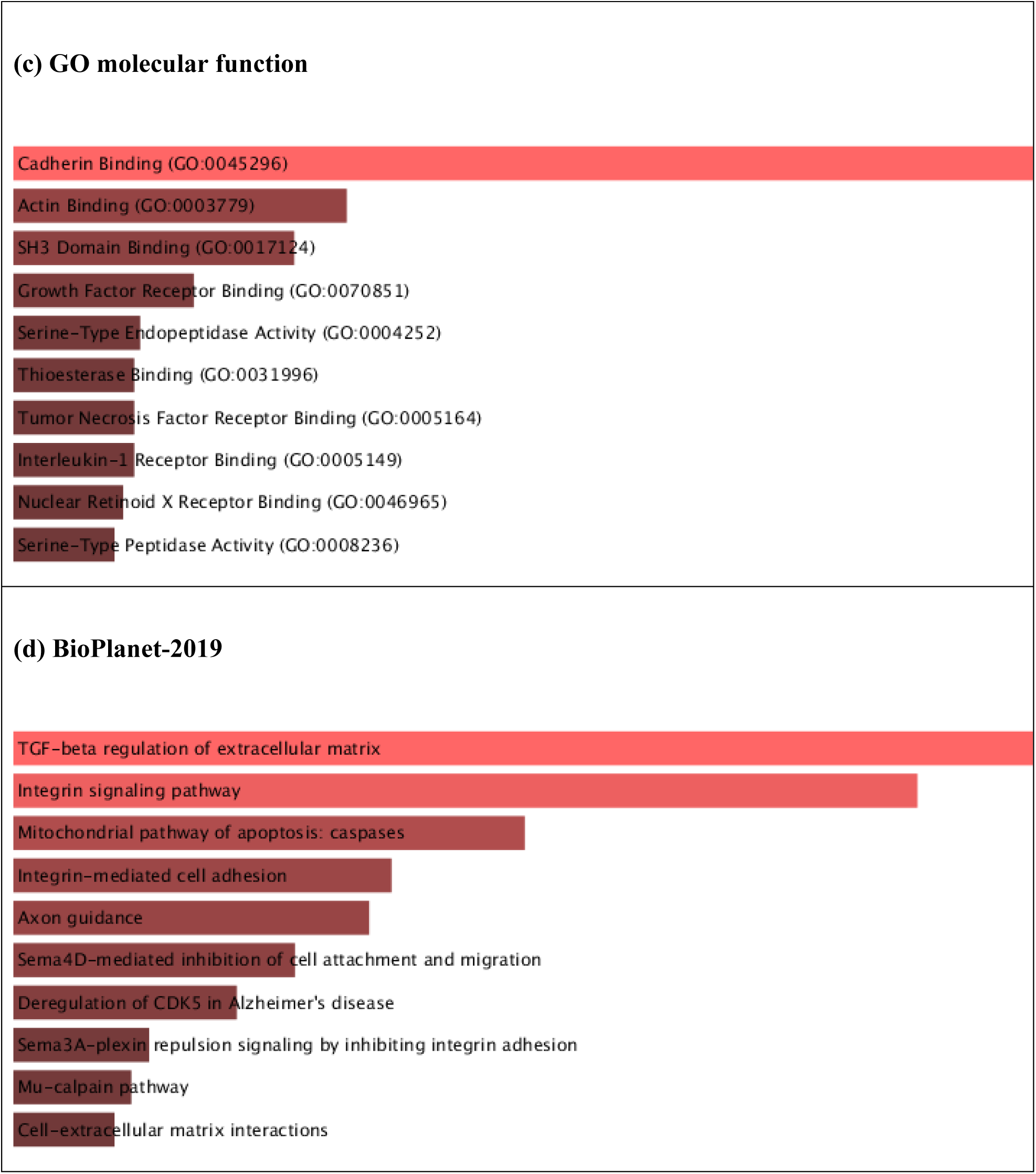
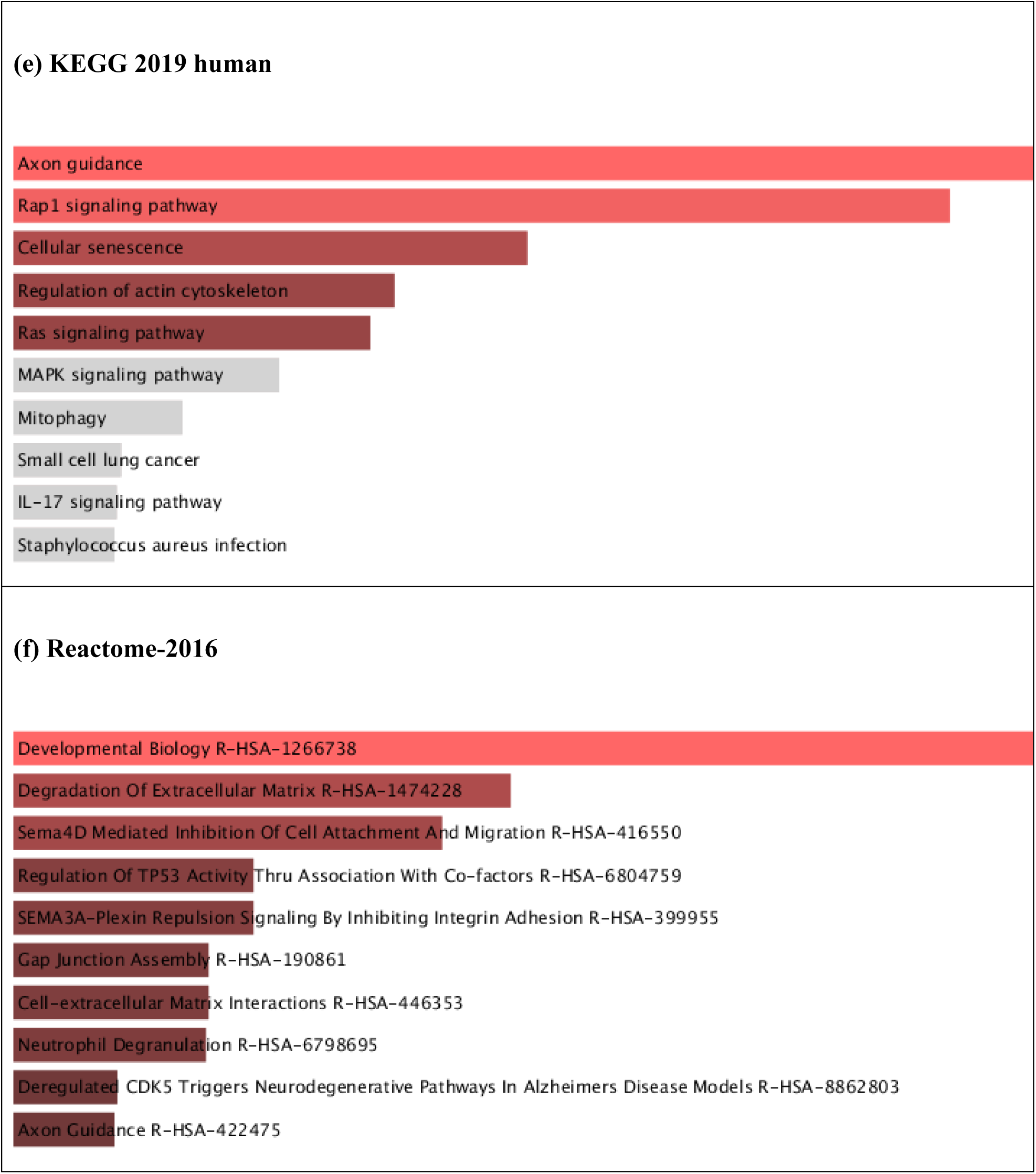
The results of the analysis of signaling pathways and gene ontology (GO). Gene ontology- (a) GO biological processes, (b) GO cellular compartment and pathway, (c) GO molecular function, (d) BioPlanet-2019, (e) KEGG-2019 human, (f) Reactome-2016 analysis of the co-expressed genes of *KCNN4* in PAAD specimens.

## 4. Conclusion

Comparing cancerous pancreatic adenocarcinoma tissues to nearby normal tissues, this study found that *KCNN4* expression is significantly higher in the former. Further analysis of abnormal promoter methylation patterns and *KCNN4* mutations may help to explain the inconsistent clinical findings of patients due to gene dysregulation with pancreatic cancer. Moreover, elevated *KCNN4* expression levels were linked to lower patient survival. It was discovered that *PPP1R13L* and other *KCNN4* genes with comparable functions were involved in processes linked to the development of pancreatic adenocarcinoma (PAAD) in pancreatic cancer tissues. When considered collectively, the data from this investigation and other research seem to support the possibility that *KCNN4* could serve as a predictive biomarker for pancreatic adenocarcinoma (PAAD). The scientific results of this investigation ought to direct future clinical research on therapeutic and diagnostic approaches based on *KCNN4* for the detection and management of pancreatic cancer. Even though there have been great advancements in the knowledge and treatment of pancreatic cancer, more research and innovation are still required given the disease’s rising prevalence and its difficult diagnostic and therapeutic cases. Reducing the global cancer burden requires addressing disparities in access to screening and treatment, enhancing early detection techniques, and creating novel, potent treatments.

